# Constitutive expression and distinct properties of IFN-epsilon protect the female reproductive tract from Zika virus infection

**DOI:** 10.1101/2022.09.01.505971

**Authors:** Rosa C. Coldbeck-Shackley, Ornella Romeo, Sarah Rosli, Linden J. Gearing, Jodee A. Gould, San S. Lim, Kylie H. Van der Hoek, Nicholas S. Eyre, Byron Shue, Sarah A. Robertson, Sonja M. Best, Michelle D. Tate, Paul J. Hertzog, Michael R. Beard

## Abstract

The immunological surveillance factors controlling vulnerability of the female reproductive tract (FRT) to sexually transmitted viral infections are not well understood. Interferon-epsilon (IFNε) is a distinct, immunoregulatory type-I IFN that is constitutively expressed by FRT epithelium and is not induced by pathogens like other antiviral IFNs α, β and λ. We show the necessity of IFNε for Zika Virus (ZIKV) protection by: increased susceptibility of IFNε^-/-^ mice; their “rescue” by intravaginal recombinant IFNε treatment and blockade of protective endogenous IFNε by neutralising antibody. Complementary studies in human FRT cell lines showed IFNε had potent anti-ZIKV activity, associated with transcriptome responses similar to IFNλ but lacking the proinflammatory gene signature of IFNα. IFNε activated STAT1/2 pathways similar to IFNα and λ that were inhibited by ZIKV-encoded non-structural (NS) proteins, but not if IFNε exposure preceded infection. This scenario is provided by the constitutive expression of endogenous IFNε. However, the IFNε expression was not inhibited by ZIKV NS proteins despite their ability to antagonise the expression of IFNβ or λ. Thus, the constitutive expression of IFNε provides cellular resistance to viral strategies of antagonism and maximises the antiviral activity of the FRT. These results show that the unique spatiotemporal properties of IFNε provides an innate immune surveillance network in the FRT that is a significant barrier to viral infection with important implications for prevention and therapy.

**Author Summary:** The female reproductive tract (FRT) is vulnerable to sexually transmitted infections and therefore a well-tuned immune surveillance system is crucial for maintaining a healthy FRT. However, our understanding of the factors that impact viral infection of the FRT and the host response are not well understood. In this work we investigate the role of a hormonally regulated type I interferon, IFN epsilon (IFNε) in control of Zika virus (ZIKV) infection of the FRT. IFNε is unique compared to other canonical type-I IFNs in that it is constitutively expressed by epithelial cells of the FRT with expression levels controlled by progesterone and not in response to viral infection. We demonstrate that IFNε has anti-ZIKV properties using a combination of IFNε KO mice, blockade of endogenous IFNε by neutralising Abs and rescue of IFNε KO mice by recombinant IFNε administered directly to the FRT. Furthermore, we complemented our in vivo studies using human FRT derived cell lines. Importantly, ZIKV NS proteins did not block IFNε expression despite their ability to antagonise the expression of IFNβ or λ. Collectively this work implicates IFNε as a key type-I IFN that provides a distinct homeostatic antiviral environment in the FRT.

## Introduction

Zika virus (ZIKV) is a mosquito-borne *flavivirus* that can also be transmitted sexually [1], and *in utero* [2] leading to foetal infection and congenital sequelae in neonates [3] including microcephaly, intrauterine growth restriction, as well as ocular and cognitive impairment [4, 5].

The innate immune response to ZIKV is critical for controlling infection [6] particularly at the mucosa of the female reproductive tract (FRT). Type-I and type-III IFNs are the body’s premier antiviral cytokines that protect against viral infection at mucosal surfaces [7]. Type-I and III IFNs orchestrate the cellular antiviral response via binding cognate receptors, IFNAR1/2 or IFNLR1/IL10Rβ, respectively. Receptor binding activates JAK/STAT signalling (as reviewed in [8]) leading to activation of thousands of Interferon Stimulated Genes (ISGs) [9]. These ISGs encode effector proteins including Viperin, IFITM family, ISG15 and IFI6 that directly inhibit ZIKV [10–13], or those with immune regulatory function [6]. Importantly, ZIKV is exquisitely sensitive to the biological effect of type-I and III IFNs evidenced by its enhanced replication in IFN receptor knockout mouse models [14, 15], but these studies did not elucidate which of the 20 IFN ligands were important. Typical type-I and III IFNs are only produced after detection of viral infection by pattern recognition receptors (PRRs) such as by the cytosolic PRR retinoic acid-inducible gene-I RIG-I [16, 17]. This delay between pathogen detection and establishment of the IFN-mediated antiviral state is exploited by ZIKV that, after entering the cell, translates non-structural (NS) proteins that can inhibit IFN production and action pathways to promote infection [18].

Unlike the typical type-I and type III IFNs, that are for the most part only expressed following viral detection, IFN-epsilon (IFNε) is a unique type-I IFN that it is expressed constitutively, independent of pathogen detection pathways and instead is regulated by female sex hormones [19]. IFNε is produced primarily by the mucosal epithelium of the FRT in both mice and humans [19, 20] and like other type-I IFNs, signals via IFNAR1/2 to induce expression of ISGs [19, 21]. This constitutive expression is important for protective mucosal immune responses to bacterial (chlamydia) and viral (HSV2) infection of the FRT [19]. Additionally, IFNε demonstrates *in vitro* activity that inhibits multiple steps of the HIV lifecycle [22]. Our current knowledge of the role of IFNs in protection from ZIKV infection of the FRT has been elegantly investigated largely by the use of IFNAR1 [23] and IFNLR1 [15] null mice that are deficient for all type-I and type-III IFN signalling, respectively. However, the particular ligands responsible for protection of the ZIKV infected FRT through these receptors are not well characterised. In this study we demonstrate the importance of IFNε as a major effector in resistance of the FRT to ZIKV infection.

## Results

### ZIKV replication is inhibited by endogenous IFNε in a mouse model of vaginal transmission

To determine the contribution of endogenous IFNε expression in the FRT, relative to other (conventional) type-I IFNs to prevent ZIKV infection, we compared the outcomes of intravaginal (iVag) infection [19] of wildtype (WT), IFNε^-/-^ or IFNAR1^-/-^ mice with 5 × 10^5^ FFU of ZIKV Puerto Rican_stain PRVABC59 (Fig. 1a).

**Figure 1:**
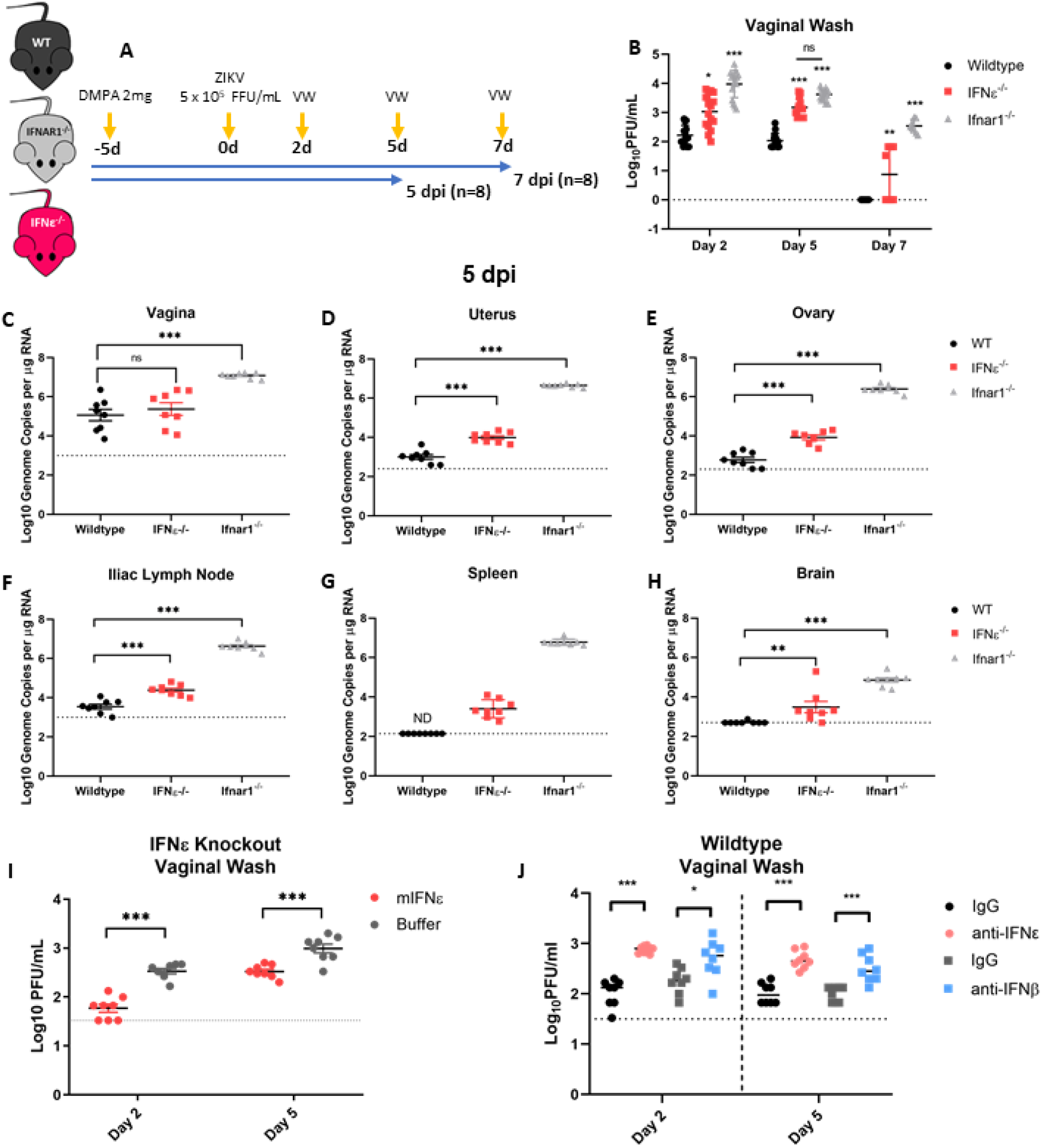
ZIKV replication is inhibited by IFNε in a mouse model of vaginal transmission. A) Experimental time line of WT (black), mice lacking IFNε (IFNε-/-) (red) or mice lacking the type-I IFN receptor (IFNAR1-/-) (grey) were infected with ZIKV at 5 × 10^5^ FFU iVag 5 days post DMPA treatment and vaginal washes were taken at 2, 5, 7 dpi. Groups of 8 mice were culled at 5 dpi and 8 were culled at 7 dpi. B) Infectious virus was measured from vaginal washes by plaque assay at 2, 5, 7 dpi. C, D, E, F, G & H) Tissues taken at 5 dpi were used to harvest RNA for analysis of viral RNA by qRT-PCR in the vagina, uterus, ovary, illiac lymph node spleen and brain respectively. I) IFNε^-/-^ mice were treated for 6h with either mIFNε or buffer prior to iVag infection with ZIKV, Infectious virus was measured from vaginal washes by plaque assay at 2 and 5 dpi. J) IFNε-/-mice were treated iVag with either mIFNε, mIFNλ2 or buffer for 6h prior to harvesting FRT tissues to determine the level of ISGs by qRT-PCR. K) WT mice mice were treated for 6h with either IFNε neutralising antibody (α-IFNε) or an isotype control (IgG) prior to iVag infection with ZIKV, Infectious virus was measured from vaginal washes by plaque assay at 2 and 5 dpi. J) WT mice were treated for 6h with either an IFNε neutralising antibody (α-IFNε), an IFNβ (α-IFNβ) neutralising antibody, or a matched isotype control prior to iVag infection with ZIKV. Infectious virus was measured from vaginal washes by plaque assay at 2 and 5 dpi. Data are presented as the mean +/− S.E.M.

In the absence of IFNε, mice were more susceptible to ZIKV infection in tissues of the reproductive tract with higher viral titres in vaginal washes (VW) 2, 5 and 7 days post-infection (dpi) (Fig. 1b). Interestingly, the levels of infectious virus in VW at 5 dpi of IFNε^-/-^ mice were not significantly different to that of IFNAR1^-/-^ mice. Higher viral load in VW correlated with increased viral RNA (vRNA) detected by qRT-PCR in both the uterus and ovary on 5 (Fig. 1d & e) and 7 dpi (Sup. 1). However, in the vagina that generally had greater virus burden (5.06 Log10) than either the uterus (3.19 Log_10_) or ovary (2.28 Log10) in WT mice, no significant difference was detected between WT and IFNε^-/-^ mice (Fig. 1c). The vagina had apparently lower IFNε RNA levels than the uterus and ovary, but this was not statistically significant (Sup. 2). Additionally, in-situ hybridisation (ISH) performed on FRT tissues at 5 dpi (Supl. 3) showed ZIKV RNA was detected in the lower FRT (LFRT) of WT, IFNε^-/-^ and IFNAR1^-/-^ mice, however this was below the limit of detection for ISH in the upper FRT (UFRT) except for IFNAR1^-/-^ mice. However, the absence of IFNε led to greater viral load in the draining illiac lymph node and spleen by 5 dpi compared to WT mice (Fig. 1f & g & h). This was despite these tissues having no detectable or very low levels of IFNε mRNA (Sup. 2) suggesting that higher viral loads may have resulted from increased dissemination from the FRT. Similar to previous reports [20, 24], we found detectable levels of IFNε mRNA in the brain of WT mice, and in the absence of IFNε, mice displayed greater levels of infection in the brain by 5 dpi. However, unlike the uterus and ovary that remained infected in the absence of IFNε, by 7 dpi the levels of ZIKV RNA detected in the lymph node, spleen and brain were equivalent to WT (Sup. 1). In contrast, ZIKV replication was higher in all tissues from IFNAR1^-/-^ mice at 5 and 7 dpi. These results suggest a temporal and specific role of IFNε in limiting ZIKV replication in the FRT and dissemination to peripheral tissues. However, additional type-I IFNs are required to control virus replication in tissues overtime.

To further characterise the role of IFNε in control of viral infection of the FRT, IFNε^-/-^ mice were reconstituted with intravaginal treatments of recombinant mIFNε or buffer alone 6 h prior to infection with ZIKV. As expected, mice treated with mIFNε had lower levels of infectious ZIKV in VW compared to buffer treated controls (Fig. 1i). To further characterise this response, we treated uninfected IFNε^-/-^ mice with recombinant mIFNε or mIFNλ-2 (4μg) and assessed antiviral ISG expression in the LFRT (Fig. 1J) and UFRT (Sup. 4) 6 h post-treatment. ISG induction occurred in response to both recombinant mIFNε and mIFNλ-2 in the vagina but not uterus or ovary, consistent with the antiviral effect of intravaginal recombinant IFN treatment being localised to the site of inoculation.

To further demonstrate the protective effect of IFNε, WT mice were injected with an IFNε blocking antibody which led to increased viral titres in comparison to the isotype control Fig. 1J). To compare the relative effect of IFNε with another type I IFN, mice were injected with an IFNβ blocking antibody and while this resulted in increased viral titres this was not as significant for that of IFNε blocking Ab at least early following infection. This effect of blocking IFNε function was evident in vaginal samples (Fig. 1J), but not in samples from the UFRT, nor distant organs (Sup. 5). In the UFRT mice controlled the infection in part due to competent IFN signalling in these tissues [25], also likely indicating the instilled IFN or antibody treatments did not reach the UFRT.

Together, this data demonstrates endogenous IFNε expression in the FRT has a significant impact on local ZIKV infection and viral dissemination at early times post infection <u>by the</u> nature of its constitutive expression prior to infection. This effect is added to by other type-I IFNs controlling infection especially at later timepoints post infection.

### Antiviral effect of IFNε on human cells of vaginal and cervical origin

Having demonstrated that IFNε plays an important role in control of ZIKV infection in mice it was important to determine the effect and mechanism of action in human cells. Therefore, we tested vaginal and cervical epithelial cells as the first cells infected by sexually transmitted pathogens [26].

These cell types were permissive to infection as evidenced by inoculation of primary transformed ectocervical (Ect1) or vaginal keratinocyte (VK2) cells with increasing MOI of ZIKV (PRVABC59). Both cell types were positive for ZIKV E-antigen by immunofluorescence staining (Fig. 2a) and released infectious virus measured by plaque assay (Fig 2b & c). We then evaluated the impact of recombinant IFNε on ZIKV infection in these cells, compared to other IFNs that are important modulators of antiviral activity at mucosal surfaces (IFNα and IFNλ-III) [8]. IFNε treatment inhibited viral infection of both Ect1 and VK2 cells by ~90%, whether determined by plaque assay or viral RNA (Ect1 vRNA: 88% reduced) (Fig 2b-e). Similar reduction in viral titres or vRNA were elicited by IFNα (Ect1 vRNA: 78%) and IFNλ (Ect1 vRNA: 83%) (Fig 2b-g), indicating that IFNε can generate an antiviral state equivalent to that of IFNα or IFNλ-III in epithelial FRT cell lines.

**Figure 2:**
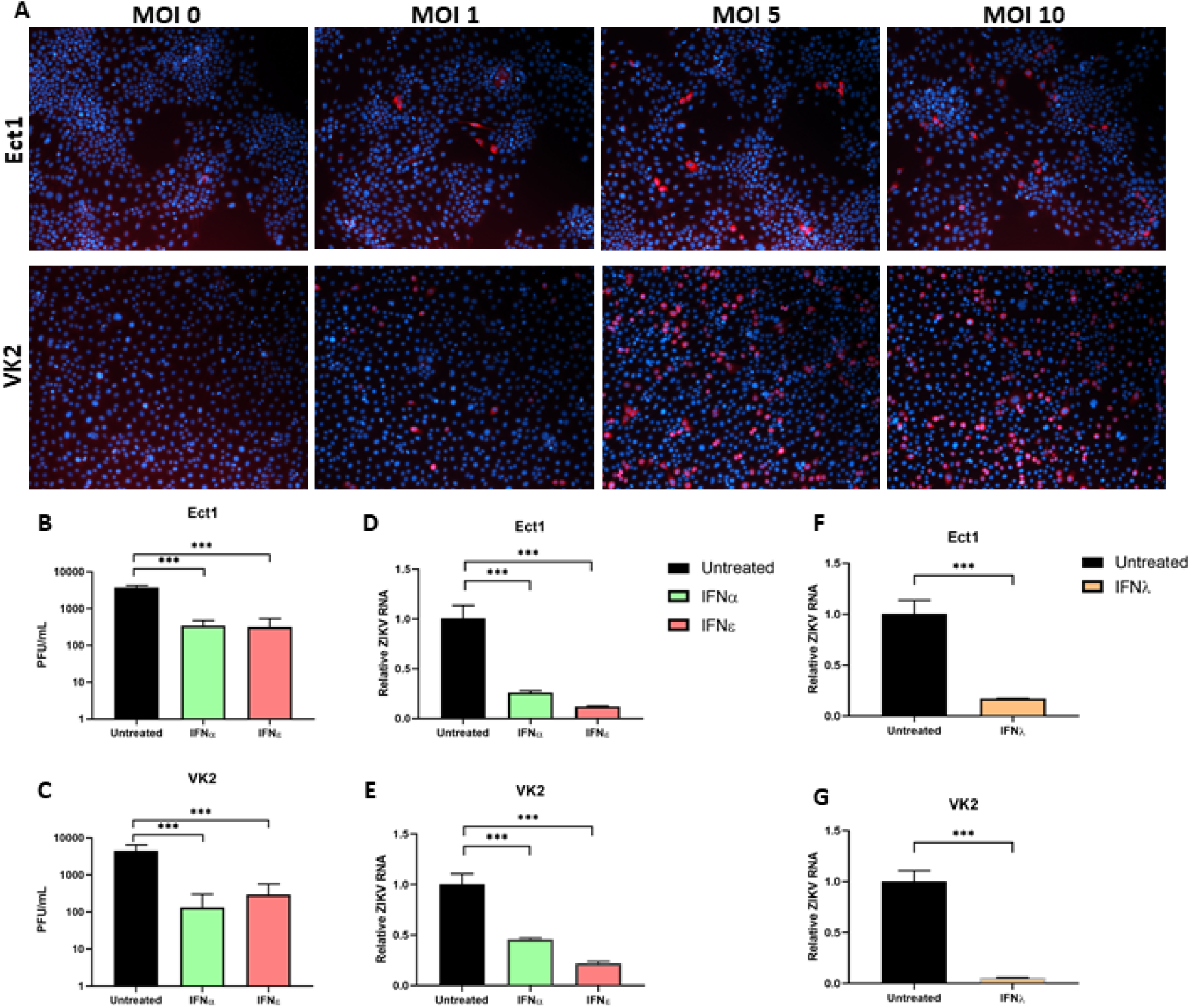
Ectocervical and Vaginal cell lines are permissive to ZIKV infection and treatment with IFNε is antiviral. A) Ect1 or VK2 cells were infected with ZIKV PRVABC59 at the indicated MOI for 24h prior to staining, anti-flavi E staining (red), DAPI (blue). A-D) Ect1 or VK2 cells were treated overnight (<u>16 hr</u>) with 100 U/mL rhIFNε, rhIFNα-2A then infected with MOI 10 or 5 respectively for a further 48h prior to collecting supernatant and RNA for plaque assay (B & C) detection of infectious virus or qRT-PCR detection (D & E)) of vRNA. F-G) Ect1 or VK2 cells were treated overnight with 100 ng/mL rhFNλ-III then infected with MOI 10 or 5 respectively for a further 48h prior to harvesting RNA for detection of vRNA by qRT-PCR. Statistical analyses are performed by one-way ANOVA compared to the untreated control (B-E) or by two tailed t-test (F & G), (n.s non-significant, * P < 0.05, ** P < 0.01). Data are presented as means +/− S.D.

To further investigate their antiviral functions, we compared cellular responses to stimulation with these IFNs in FRT cells using whole transcriptome profiling by RNA-seq analysis. Interestingly, IFNε and IFNλ-III had almost indistinguishable gene signatures at this time point (Fig. 3a, Sup. 6) which contrasts recent reports comparing type-I, IFNβ, and type-III IFNs that found several differently regulated ISGs [15, 27]. As expected, all IFN treatments significantly upregulated canonical ISGs in both cell types compared to untreated controls (Fig 3b, c & d). Consistent with the reported specific activities of these IFNs [21, 28, 29], IFNα induced the strongest response, inducing greater numbers and magnitude changes of responsive genes (Fig 3b).

**Figure 3:**
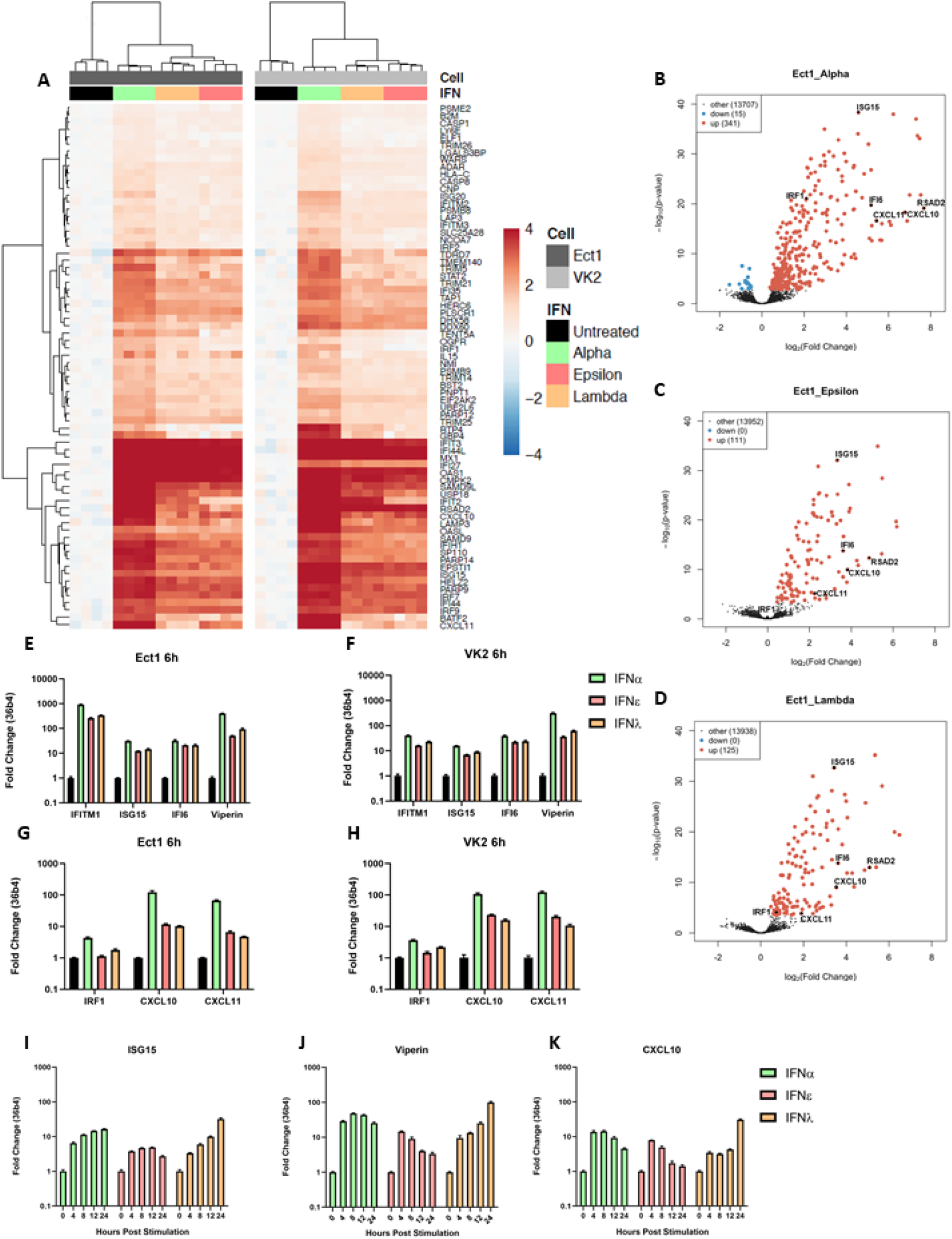
IFNε displays typical type-I IFN kinetics but induces an antiviral gene signature like IFNλ-3 at early time points in ectocervical and vaginal cells. Ect1 and VK2 cells were treated with IFNε, IFNα-2a or IFNλ-3 (100 ng/mL) or left untreated (n = 4) for 6hr prior to RNA-seq analysis (NextSeq550 V2.5). Differentially expressed genes were determined with a 1.2-fold cut-off and adjusted p-value < 0.05. A) Heat map showing expression of key ISGs. B, C & D) Volcano plots indicating up or downregulation of genes after IFNα-2a, IFNε or IFNλ-3 treatment, respectively. E & F) Confirmation of anti-ZIKV ISGs (ISG15, IFI6, IFITM1, Viperin) expression by qRT-PCR. G & H) Confirmation of pro-inflammatory ISGs (IRF1, CXCL10, CXCL11) expression by qRT-PCR. I, J & K) Ect1 cells were treated with IFNε, IFNα-2a or IFNλ-3 (100 ng/mL) and RNA was harvested from untreated cells (t = 0) and at the indicated time points post stimulation. qRT-PCR was performed to detect expression of ISG15, Viperin and CXCL10.

Next, we performed qRT-PCR analysis to confirm expression of key antiviral (Fig. 3e & f) or proinflammatory (Fig 3g & h) genes. Key ISGs that protect against ZIKV infection (Viperin [10], ISG15 [30], IFITM1[11], IFI6 [13]) were upregulated in all three treatments. We also noted that while both type-I and type-III IFNs induced expression of the proinflammatory CXCR3 ligands, CXCL10 and CXCL11, and that their expression was significantly lower in IFNε and IFNλ treatments. Interestingly, a recent report has suggested that temporal ISG induction by type-I and III IFNs provides a collaborative antiviral response, with type-I IFNβ promoting inflammation via an IRF-1 dependant inflammatory response. Consistent with this observation we also noted an increase in IRF-1 expression in Ect1 and VK2 cells stimulated with IFNα as compared to stimulation with IFNε and IFNλ-III (Fig. 3a, h,i). Taken together, these gene expression profiles suggest that IFNα, like IFNβ, drives a greater proinflammatory phenotype in comparison to IFNε and IFNλ-III.

To further understand the spaciotemporal action of IFNε, we compared ISG induction in Ect1 cells responding to IFNε, IFNα-2A, and IFNλ-III treatments over time. IFNε induced an early but transient induction of ISGs (ISG15, Viperin, CXCL10) while IFNα induced early induction of ISGs that was maintained over the 24 h time course (Fig. 3i, j & k) and IFNλ a later, gradual induction of ISGs over 24 hrs as seen previously in liver and lung epithelial cell lines [27, 31–35]. Collectively these transcriptional signatures suggest that these IFNs have evolved to perform specialised functions to coordinate antiviral responses spatiotemporally to combat viral infection of the FRT mucosa. Of these functions the apparent ability of IFNε to induce lower levels of IRF1 and proinflammatory genes in the FRT, like IFNλ rather than IFNα, warrants further investigation.

### Treatment of cells with IFNε prior to infection precedes ZIKV evasion of type-I and type-III IFN signalling pathways

To understand the importance of IFNε’s constitutive expression on ZIKV evasion of IFN responses in humans, compared to IFN expression that is induced following viral infection, we treated the placental trophoblast cell line HTR8, with IFNε or IFNα either pre- or post-infection with ZIKV (Fig. 4a). These cells were chosen as they infect to higher level compared to the VK2 and Ect1 cells and they mimic target cells for ZIKV transplacental transmission at the foetal-maternal interface [36]. Pre-treatment with IFNs imparted a strong block to infection whereas treatment at 3 hpi was less effective and antiviral effects were almost lost when treatment was delayed until 24 hpi for both IFNε and IFNα (Fig. 4 b&c). This observation is consistent with similar studies for other flaviviruses, showing a rapid shut-down of IFN signalling following viral entry and detectable NS protein expression as early as 3 hpi [37, 38]. As expected, this effect was also observed for IFNλ-III, since this IFN relies on an overlapping signalling pathway with type-I IFNs (Fig. 4d & e).

**Figure 4:**
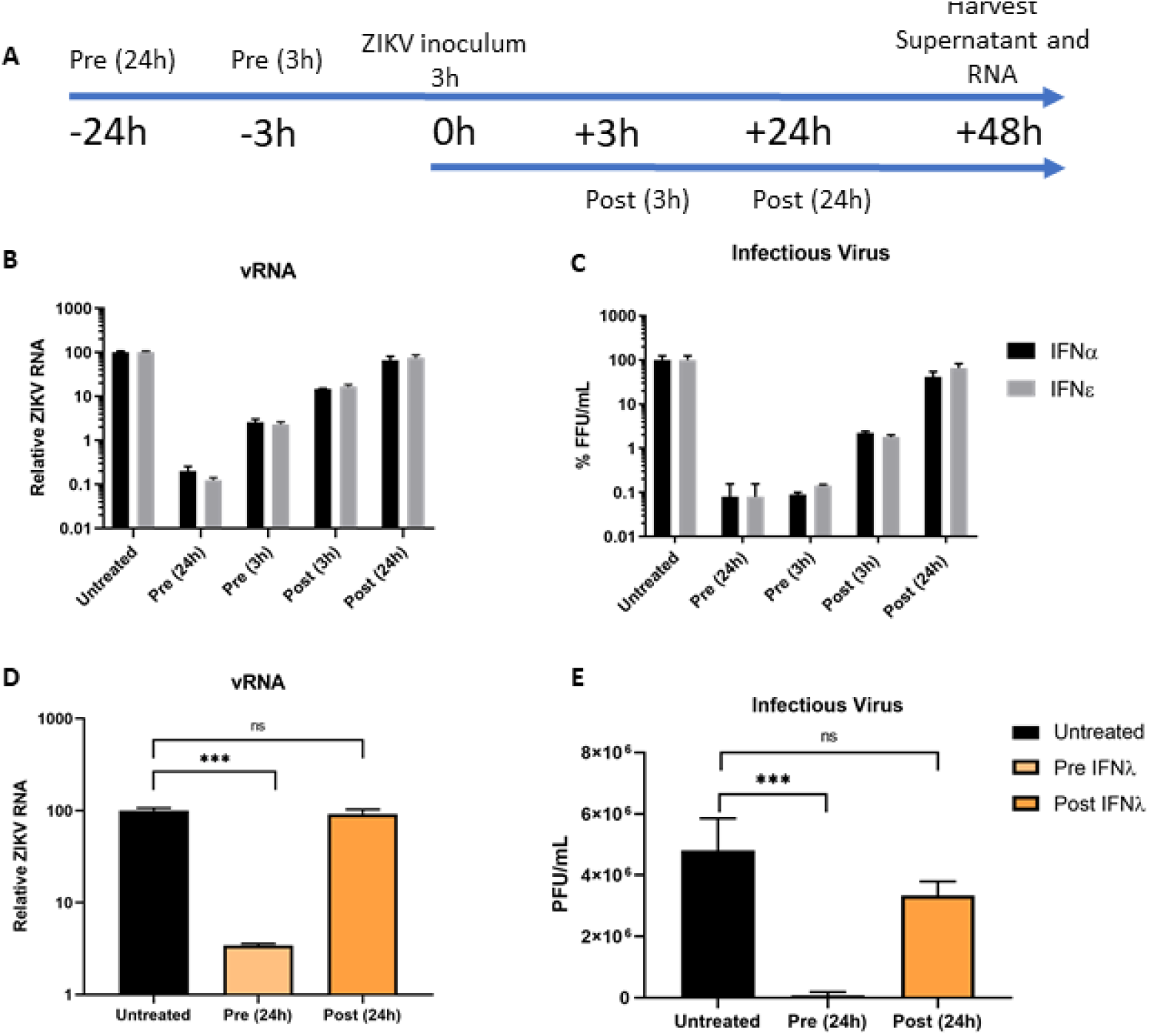
ZIKV evades type-I and III IFN antiviral activity post-infection. A) Timeline for IFN treatment regimes in HTR8 cells. B & C) HTR8 cells were infected with ZIKV PRVABC59 at a MOI of 1, cells were either primed with the indicated IFN (mIFNε 10 U/mL or hIFNα-2A 500 U/mL) or treated post infection. 48 hpi supernatants and RNA were harvested for determination of infectious virus by Focus Forming Assay or quantification of viral RNA by qRT-PCR. D & E) HTR8 cells were infected with ZIKV PRVABC59 at a MOI of 1, cells were either primed for 24h with hIFN λ-III (100 ng/mL) or treated post 24h post infection. Viral RNA and supernatant were collected 48hpi for determination of infectious virus by Focus Forming Assay or quantification of viral RNA by qRT-PCR. Statistical analyses are performed by or one-way ANOVA compared to the untreated control, (n.s non-significant, * P < 0.05, ** P < 0.01). Data are presented as means +/− S.D.

To determine the step in the signalling cascade antagonised by ZIKV, we investigated STAT1/2 nuclear translocation in response to IFNε by immunofluorescence <u>in VK2 and Ect1 cells</u>, and STAT1/2 phosphorylation 30 min post-treatment by western blotting in HeLa cells infected with ZIKV. HeLa cells were chosen for immunoblot experiments as the low level of infection in VK2 and Ect 1 confounded interpretation of the data due to the bystander effect from uninfected cells. ZIKV infected VK2 (and Ect1, HeLa cells supplementary) cells had reduced nuclear translocation of both STAT1 and STAT2 (Fig 5a, b). This nuclear translocation was also noted in HeLa cells with an associated reduction in STAT1/2 phosphorylation (Fig 5c). However, STAT2 translocation, phosphorylation and total protein levels were more potently reduced by ZIKV infection compared to STAT1. A similar inhibition of STAT1 and STAT2 function was observed following stimulation of infected cells with IFNα (Fig. 5d), suggesting a common mechanism of ZIKV-mediated evasion was responsible for inhibiting the pathway downstream of both IFNs.

**Figure 5:**
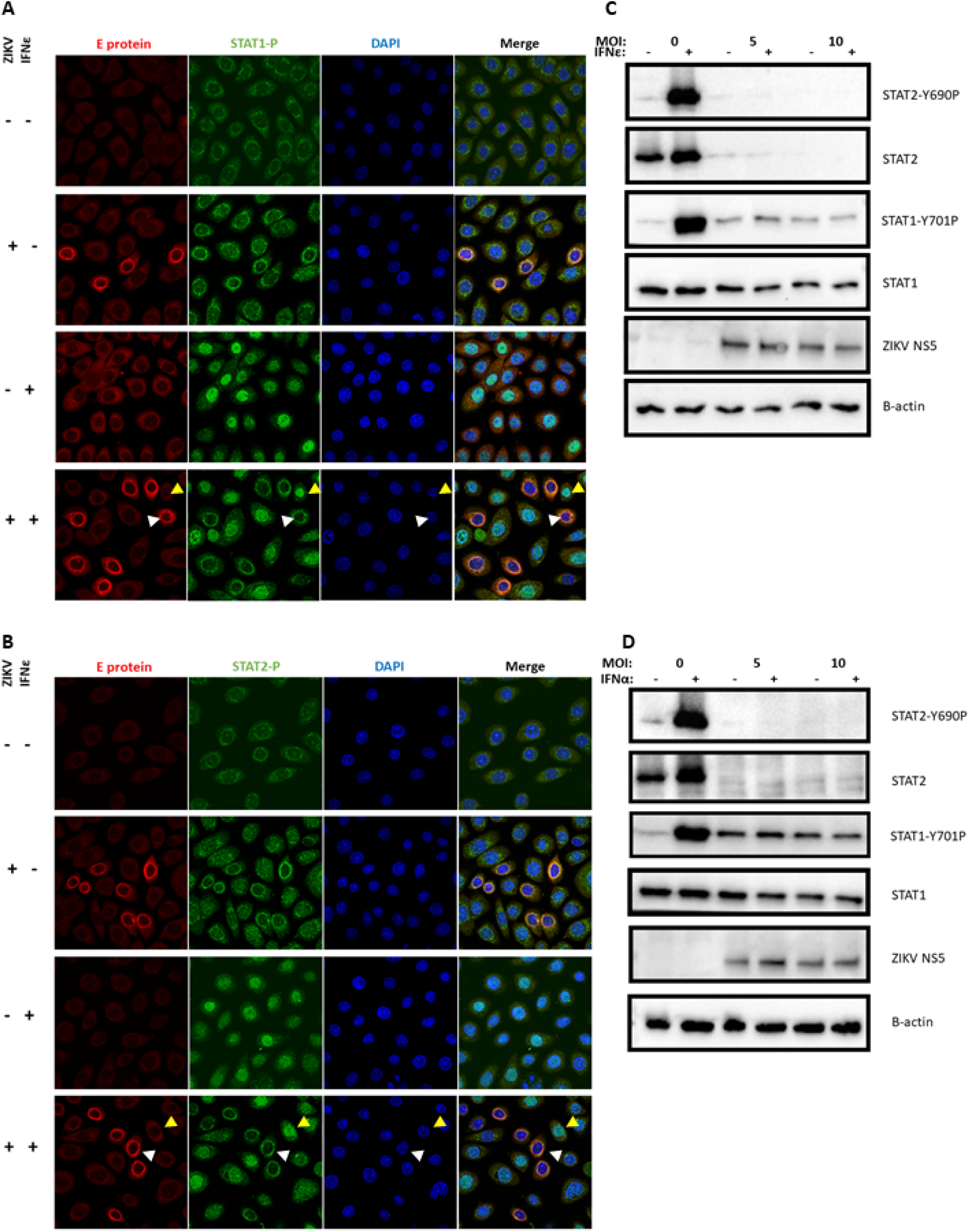
Antiviral protection mediated by IFNε and other type-I IFNs is potently inhibited due to ZIKV inhibition of STAT1/2 signalling. A & B) Ect1 cells were infected with ZIKV MOI of 10, 24 h post infection cells were stimulated with hIFNε (100 U/mL) for 30 min then fixed with acetone/methanol for detection of ZIKV E-antigen (red) and phosphorylated STAT1 or STAT2 proteins (green) by indirect immunofluorescence, DAPI (blue), infected cells (white arrows) and indicate uninfected bystanders (gold arrows). C & D) HeLa cells were infected at the indicated MOI of ZIKV 24h prior to stimulation with the either mIFNε (10 U/mL) or hIFNα-2A (500 U/mL) for 30 min, lysates were harvested for immunoblot of STAT1/2 protein and phosphorylated STAT1/2 proteins.

Next, we determined if inhibition of STAT2 activation by IFNε was driven by the viral determinants previously described for ZIKV antagonism of IFNα [39]. HeLa cells were again used to overcome the refractory nature of VK2 and Ect1 cells to transfection. Cells transfected with plasmids expressing ZIKV NS5, or an empty vector were stimulated with IFNε or IFNα and 30 minutes post stimulation, STAT1/STAT2 activation was assessed as above. In the presence of NS5 protein, total STAT2 protein was reduced with a concomitant decrease in STAT2 phosphorylation in response to IFNε (Fig. 6a) and IFNα (Fig. 6b). Marginal, if any, impact of NS5 was observed on STAT1 phosphorylation (Fig. 6a,b). Interestingly, total STAT1 and phosphorylated STAT1 in the presence of ZIKV NS2B/3 and IFNs remained constant despite previous findings that ectopic expression of NS2B/3 protein inhibits STAT1 activation [40]. To confirm the impact of reduced STAT2 phosphorylation on downstream signalling we performed a dual luciferase assay for ISRE activity and qRT-PCR for ISGs. Accordingly, NS5 expression also inhibited downstream ISRE promoter activity (Fig. 6c) and ISG expression (representative gene ISG15 shown for simplicity) in HeLa cells in a dose dependent fashion (Fig. 6d).

**Figure 6:**
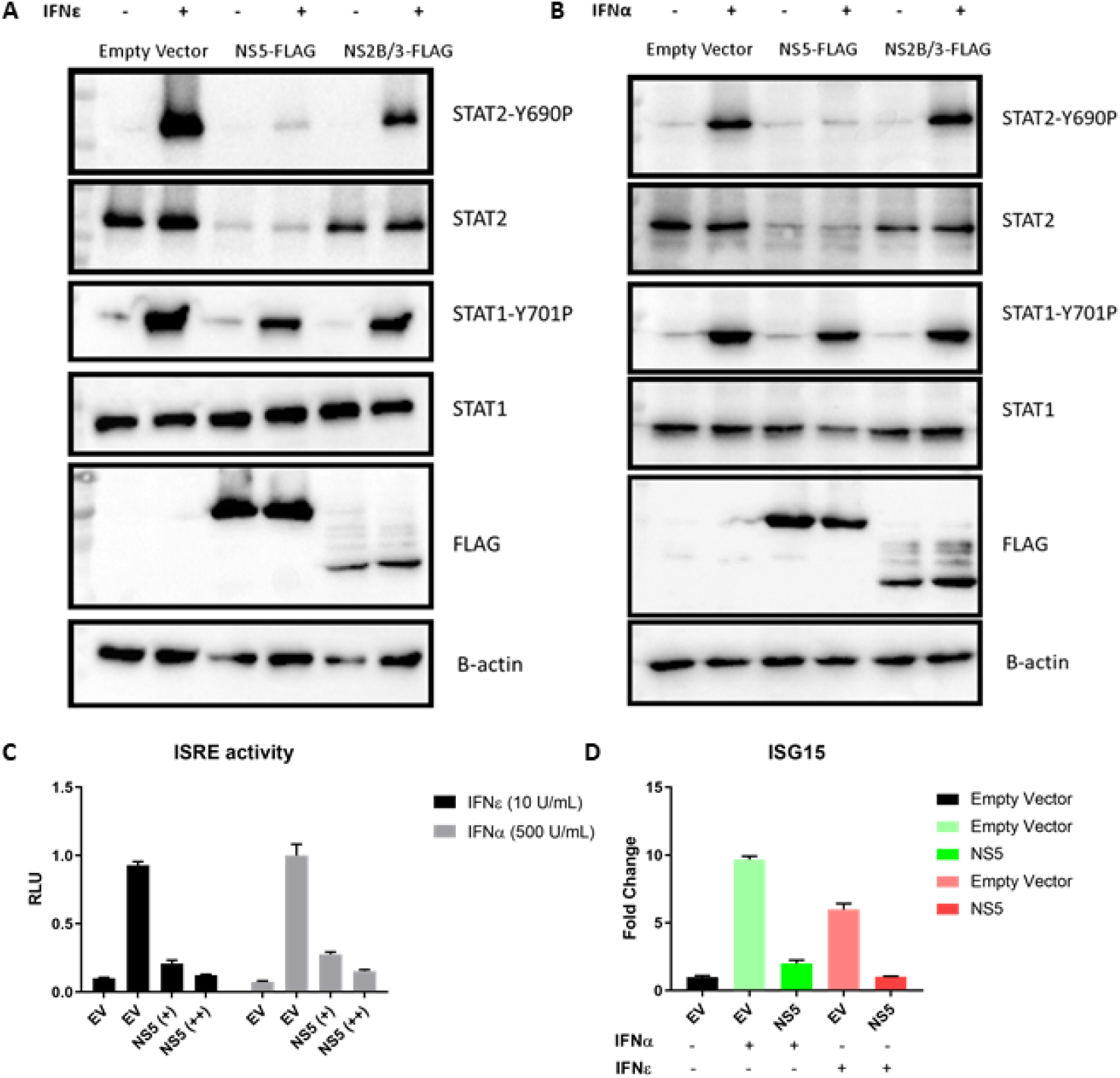
Evasion of IFNε antiviral activity is mediated by ZIKV NS5 degradation of STAT2. A & B) HeLa cells were transfected with pCDNA6.2-ZIKV-NS5-FLAG, NS2B/3-FLAG or empty vector control and 24 h later stimulated for 30min with the indicated IFN prior to assessing total and phosphorylated STAT1/2 by immunoblotting. C) ISRE promoter activity was assessed following 24 h IFNε stimulation in the presence or absence of ZIKV NS5A expression. D) HeLa cells were transfected with pCDNA-NS5-FLAG expression plasmid or an empty vector control (EV), 24h post transfection cells were stimulated with the indicated type-I IFN for 6 h prior to harvesting RNA for qRT-PCR analysis of ISG15 expression.

Taken together our results demonstrate the potent ability of ZIKV to shut-down JAK-STAT signalling by conventional (IFNα) and the unusual (IFNε) type I IFNs. ZIKV NS5 induced STAT2 degradation is likely to impact type III as well as type I IFNs shown above. This highlights the importance of priming with IFN before infection is established to limit virus spread and therefore indicates the likely significance of IFNε as the only constitutively expressed IFN detectable in the FRT [41] providing antiviral protection prior to expression of viral antagonists of IFN signalling.

### IFNEε constitutive expression is not inhibited by ZIKV infection or NS proteins

The constitutive expression of IFNε in the FRT in the absence of viral infection is hypothesised to circumvent pathogen-mediated IFN evasion helping to protect the FRT from viral and bacterial infections [42]. However, this assumes infection cannot inhibit IFNε expression, therefore we asked if ZIKV infection could antagonise the expression of IFNε.

HeLa cells are known to express IFNε RNA at levels similar to FRT cell lines [19], therefore we used these to investigate expression of IFNε compared to other type-I and III IFNs in response to ZIKV infection or poly I:C stimulation (viral mimic). At a basal level, IFNε was highly expressed compared to other type-I and III IFNs (Fig. 7a). Consistent with previous reports [19], IFNε was not significantly induced in response to either ZIKV infection or poly I:C transfection. Conversely, PRR-mediated induction of type-I IFNβ and type-III IFNλ was required to generate expression levels equivalent to basal IFNε in these cells. Interestingly, we did not see ZIKV mediated dampening of poly I:C induced IFNβ induction when cells were infected prior to poly I:C stimulation as reported previously by our group [10] and others [40]. However, previous experiments were conducted in either Huh-7 or A549 cells, suggesting the PRR pathways that contribute to HeLa cell IFN expression may differ and are not as efficiently inhibited by ZIKV.

**Figure 7:**
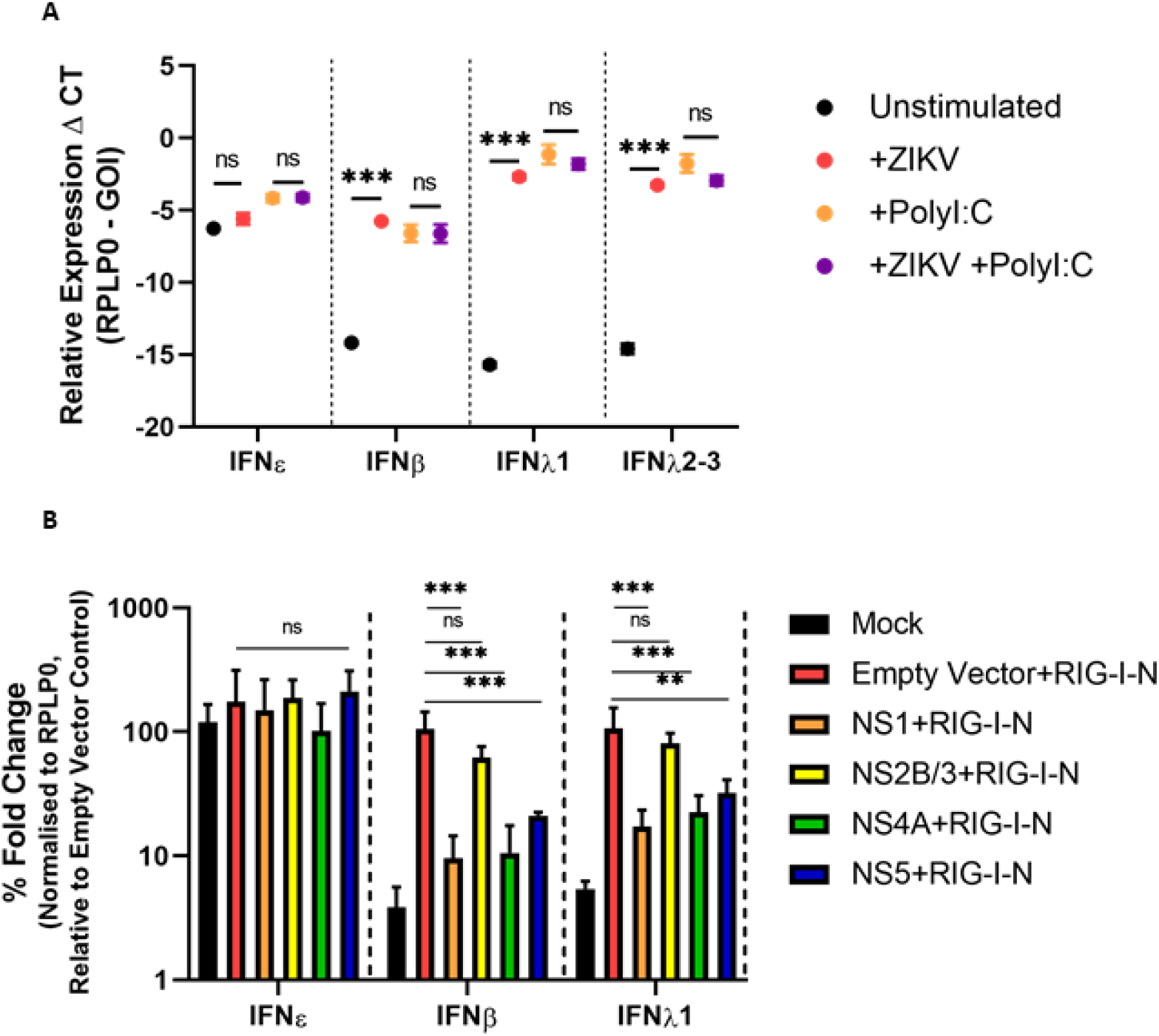
IFNε RNA expression is not reduced by ZIKV infection or NS protein expression. A) HeLa cells infected for 16h MOI 1 ZIKV PRV prior to stimulation with Poly I:C 1 μg for 8 h prior to assessing IFN gene expression by qRT-PCR. B) HeLa cells were co-transfected with expression plasmids encoding ZIKV NS1/2B/3/4A/5 or empty vector (pCDNA6.2) and the RIG-I-N plasmid or were mock transfected (Lipofectamine only), 24 h later RNA was harvested prior to assessing induction of type-I and type-III IFNs by qRT-PCR. Statistical analyses are performed by two-way ANOVA (A) or one-way ANOVA compared to the empty vector control (B), (n.s non-significant, * P < 0.05, ** P < 0.01). Data are presented as means +/− S.D.

To focus on the main pathway responsible for inducing IFN expression in response to ZIKV infection (RIG-I/MAVS) [17, 23] we co-transfected HeLa cells with a plasmid expressing constitutively active RIG-I (RIG-I-N [43]) and either vectors expressing ZIKV-NS1/NS4A/NS2B-3/NS5-FLAG or an empty vector control (see Sup. 7 for protein expression levels). These NS proteins are known to restrict the expression of other type-I IFNs in response to viral infection [44–48]. Expression of RIG-I-N alone resulted in the upregulation of IFNβ and type-III IFNλ and this was supressed by co-expression of ZIKV-NS1, NS4A, or NS5 but not NS2B/3 (Fig. 7b). Conversely, constitutive expression of IFNε RNA was unaffected by RIG-I pathway stimulation or by the presence of NS proteins in these cells.

Together this data demonstrates IFNε constitutive expression is not altered following ZIKV infection. This contrasts with other type-I and III IFNs (IFNβ and IFNλ-I) that rely on PRR activation for their expression and are therefore susceptible to inhibition by ZIKV NS proteins.

## Discussion

ZIKV is the only *flavivirus* known to transmit both sexually and *in utero*, making it a unique threat to the FRT immunological niche, both during pregnancy and under normal physiological conditions. Here we demonstrate the non-redundant role of a relatively uncharacterised type-I IFN; IFNε, in protecting the FRT from ZIKV infections both *in vitro* and in a mouse model of vaginal infection.

To assess the contribution of endogenous IFNε in controlling ZIKV FRT infections we used a murine model of vaginal infection. In this model DMPA treatment was used to synchronise mice to diestrus, rendering mice susceptible to ZIKV iVag infection as reported by other groups [15, 23, 49]. Since IFNε is hormonally regulated its expression is reduced during diestrus and pregnancy in mice [19]. This reduction in IFNε may contribute to the natural susceptibility of WT mice during diestrus [23, 49] and pregnancy [23] to iVag ZIKV infection compared to systemic inoculation methods. Importantly, this means our data must be viewed as “no” versus “low” IFNε in the FRT when comparing IFNε^-/-^ to WT mice. ZIKV infection via iVag inoculation was compared between WT, IFNε^-/-^ and IFNAR1^-/-^ mice.

Although DMPA treated WT mice expressed low levels of IFNε, we found that its presence in the FRT was sufficient to offer significant protection against ZIKV iVag infection compared to mice lacking IFNε entirely. This protection was most significant during the early stages of infection as indicated by the high levels of infectious virus recovered in the VW of IFNε^-/-^ mice that closely mirrored levels observed in the highly susceptible IFNAR1^-/-^ mice. Interestingly, the antiviral activity of endogenous IFNε primarily acted in the UFRT tissues and not the vagina, despite the vagina expressing high levels of IFNε mRNA compared to peripheral tissues such as the spleen. Notably, there was also a smaller difference in vRNA levels between WT and IFNAR1^-/-^ mice in this tissue compared to UFRT tissues. This demonstrates a lesser overall impact of type-I IFN signalling in the vagina and agrees with recent reports showing a small impact of IFNAR1^-/-^ in ZIKV infected LFRT compared to the UFRT in mice, reportedly due to lower relative PRR expression in LFRT that would limit expression of PRR induced type-I IFNs such as IFNα and IFNβ [25]. Our data suggests the LFRT is highly susceptible to ZIKV infection independent of endogenous or induced IFN expression, potentially explaining ZIKVs aptitude for sexual transmission in humans. Comparatively we demonstrate that IFNε expressed in the UFRT, the site of *in utero* transmission, is essential for ZIKV infection control.

In apparent contrast to our observations that indicate IFNε plays a significant role in protecting the FRT from ZIKV infection, multiple groups have observed type-I and III IFN independent protection of the FRT during estrus in mice [15, 50]. This appears counterintuitive to IFNε playing a significant role since mice in estrus have the highest levels of IFNε expression in FRT tissues [19]. This mechanism of estrus-dependent, IFN-independent protection against ZIKV infection in mice is yet to be fully characterised. However, one plausible explanation is that this resistance to viral infection is mediated by thickening of the epithelial layer during estrus in both upper and lower reproductive tract of mice, presenting an additional physical barrier to infection. Comparatively in humans, the epithelium of the lower reproductive tract remains constant over the menstrual cycle [51]. This additional barrier during estrus may mask the contribution of IFNε during this stage of the cycle in mice. Therefore, it is plausible the effect of IFNε may be more significant in human sexual transmission due to physiological differences in FRT biology. Additionally, ZIKV NS5 mediated evasion of human but not mouse IFN signalling via STAT2 degradation [39] likely enhances the antiviral contribution of ZIKV-induced IFNs in WT compared to IFNAR/IFNLR^-/-^ mouse models.

Interestingly, IFNε also significantly decreased the dissemination of ZIKV to peripheral tissues in WT mice by 5 dpi. Since IFNε is primarily expressed in the FRT, these results suggest that greater replication in the FRT drives increased dissemination of the virus to the peripheral organs. However, unlike the spleen and lymph node that do not express IFNε, we detected relatively high levels of IFNε mRNA in the brain (see Sup. 1) consistent with reports by others [20, 52]. If IFNε protein is functionally expressed in the brain it may have a role in protecting against ZIKV as a neurotropic *flavivirus*. Interestingly, one study in humans has linked a polymorphism of IFNε to cerebral haemorrhage, indicating it may be functionally expressed in this tissue [53]. Studies to determine the antiviral role of IFNε in the brain would be an area of interest to ZIKV and other neurotropic viruses. Collectively this data has demonstrated that IFNε plays a significant and non-redundant role at physiological levels during early infection of the FRT as a critical barrier to preventing systemic dissemination in non-pregnant adult female mice.

Understanding the capacity to modulate immune responses is the first step towards developing new immune therapies. Here we have shown modulation of IFNε levels in the LFRT can alter the infectivity of mice by ZIKV intravaginal infection. Antibody mediated neutralisation of IFNε in WT mice significantly increased infection. On the other hand, iVag treatment with IFNε was able to restore antiviral activity in IFNε^-/-^ mice. Furthermore, we investigated the immune stimulatory properties of intravaginally administered recombinant IFNε. This produced increased ISG expression in the vagina without significantly inducing ISGs in the UFRT. This indicates short iVag IFN treatments mainly act to prevent establishment of infection in the LFRT likely protecting against ascending infection and female to male sexual transmission.

To extend these observations to human cell culture we next investigated the antiviral properties of IFNε on FRT cell lines. Pre-treatment with either IFNε, IFNα or IFNλ3 protected Ect1 and VK2 cells from ZIKV infection, indicating the importance of both type-I and III IFNs in protection of the FRT in humans. We found that IFNε generated antiviral responses in FRT Ect1 and VK2 cell lines by inducing a typical IFN antiviral signature, including upregulation of several ISGs known to inhibit ZIKV infection. Interestingly, the gene signature of IFNε treatment was almost indistinguishable to IFNλ at this time point (6 h) despite signalling through different receptors. Both IFNε and IFNλ were found to induce lower levels of the transcription factor IRF1 and proinflammatory genes compared to conventional type-I IFNα treatment at the same concentration. Confirmation by qRT-PCR of key anti-ZIKV ISGs or proinflammatory genes indicated fold induction consistent with the reported specific activities of these IFNs, with IFNα inducing a more potent response than either IFNε or IFNλ at the same ng/mL concentration. Together this data shows at early times post-stimulation, IFNε induces a typical antiviral gene signature in FRT cell lines, and like type-III IFNs may have a lower propensity to induce inflammation via CXCR3 ligand regulation compared to conventional type-I IFNs. We propose the lower specific activity of IFNε provides an evolutionary advantage because of its constitutive presence in the FRT, striking a balance between limiting infection, inflammation, and normal reproductive function in these tissues. Additionally, we found that the kinetics of gene induction in response to IFNε in FRT cell line was rapid, having highest levels of ISGs at early times and waning by 24 h. Comparatively, IFNλ displayed typical type-III induction having gradually increasing gene expression with the greatest levels of gene induction at 24 h. This suggests that type-I and type-III IFNs likely have differing spatiotemporal roles in controlling ZIKV infection and most likely other infections of the FRT.

The importance of both type-I and type-III IFNs protecting against ZIKV infection is now well established in mice [15, 23, 49, 54, 55]. However, in humans and other primates, ZIKV has adapted to evade IFN responses via multiple host-viral interactions that restrict the effectiveness of IFN-mediated protection [39, 40, 44, 56]. Understanding and exploiting this axis of host-virus interaction may improve patient outcomes. Here we have shown ZIKV can evade both type-I and III IFN mediated antiviral activities downstream of receptor binding within hours after infection is established. ZIKV was able to limit the induction of ISGs via blocking both STAT1 and STAT2 activation. The key molecular mechanism underlying this effect was found to be NS5 mediated degradation of STAT2 protein as previously described for IFNα [39] and importantly for this study IFNε and most likely IFNλ. Equivalent viral evasion highlights the similarities between the type-I and III IFN signalling pathways that contribute to anti-ZIKV activity. Furthermore, ZIKVs efficient evasion of both type-I and III IFN antiviral activity after infection is established, highlights the importance of a rapid induction of ISGs or priming cells with IFN to effectively prevent infection. This also suggests that at times when IFNε is reduced, such as early pregnancy [19, 57], the FRT is likely more vulnerable to ZIKV infection. Importantly, IFNε is the only IFN known to be produced constitutively by mucosal surfaces of the non-pregnant FRT [41] suggesting it may have a significant impact on human FRT infections.

Additionally, we characterised the constitutive nature of IFNε expression with respect to virus infection and showed that in contrast to type-I and III IFNs, endogenous expression of IFNε was not susceptible to ZIKV-mediated evasion. This demonstrates for ZIKV infections, the FRT can be pre-emptively tuned to an antiviral state and that it appears to be a unique property of IFNε, stemming from the different regulatory pathways that govern its expression. Further exploration into the stimuli and pathways that regulate IFNε expression in the human FRT will therefore be important to downstream therapeutic applications.

In summary we have shown that IFNε is an important mediator of antiviral activity in the FRT of both mice and humans. This activity is effective to reduce ZIKV iVag infection in mice and infection in FRT cell lines. The constitutive presence of IFNε in the non-pregnant FRT is likely significant in human infections as ZIKV can effectively evade post-infection IFN responses but is strongly inhibited by prophylactic treatment.

## Methods and Materials

### Virus

Low passage ZIKV strain PRVABC59 was propagated in C6/36 mosquito larvae cells. Infectious virus stock titres were determined by Focus Forming Assay on Huh7.5 cells.

### Cell lines, reagents and recombinant IFNs

HeLa and Vero E6 cells were cultured in DMEM supplemented with 10% FCS and 1% Penicillin/Streptomycin. HTR8 cells were maintained in RMPI with 10% FCS and 1% Penicillin/Streptomycin. Ect1 and VK2 cells were cultured in Keratinocyte Serum Free Media supplemented with 0.1 ng/mL human recombinant EGF, 0.05 mg/mL bovine pituitary extract, additional calcium chloride 44.1 mg/mL and 1% Penicillin/Streptomycin. All cell lines were cultured at 37 °C with 5% CO_2_. Mouse and human recombinant IFNε protein (mIFNε, hIFNε) were made in house as previously described [21, 22]. Commercial IFNs were hIFNα-2A (PeproTech), hIL-28B (R&D systems) and mIFNλ2 (PeproTech). Due to the limited supply of hIFNε, large scale *in vitro* experiments on HTR8 or HeLa cells used mIFNε (10 U/mL) and hIFNα-2A (500 U/mL). These concentrations were titered by direct comparison of receptor activation levels and antiviral properties (see Sup. 8). For specific use of antibodies see supplementary methods. IFNε neutralising monoclonal antibody was generated inhouse from immunisation of IFNε^-/-^ mice with recombinant mulFNε (Lim.S., Hertzog, PJ *et al* manuscript in preparation).

### Mice

Sexually mature female mice aged between 6 – 12 weeks at the time of infection were used in these experiments. Mice from each genetic background were age matched between experiment groups to minimise the impact of differing susceptibility. C57BL/6 (WT) mice were purchased from Monash Animal Services and acclimatised for one week prior to infection. IFNε^-/-^ and IFNAR1^-/-^ mice on C57BL/6 background were maintained in house. All mouse experiments were approved by the Monash Medical Centre B ethics committee (#MMCB/2017/04).

### Intravaginal infection and treatments of mice

All mice were treated with 2 mg Depo-ralovera DMPA (Kenral) subcutaneously 5 days prior to infection to synchronize the estrus cycle into diestrus phase on the day of infection as previously described [19]. On day 0 mice were anesthetised using isoflurane then inoculated intravaginally with 5 × 10^5^ FFU ZIKV in 10 μL PBS, then kept under aesthetic for 2-3 minutes to promote viral adherence. Mice were monitored daily to assess clinical signs of disease: 0, no apparent signs of disease; 1, genital redness or minor genital swelling; 2, genital redness and genital swelling, huddled and inactive; 3, severe lethargy and little response to handling. At day 5 or 7 mice were culled, and tissues were extracted. Tissues for RNA extraction were flash frozen in liquid nitrogen and tissues for IHC or ISH staining were fixed in 10% neutral buffered formalin for a minimum of 24 h prior to embedding in paraffin. Treatments with 4 μg mIFNε or 100 μg IFNε neutralising antibody (made in house) were administered intravaginally 6 h prior to infection (or tissue collection) and again on day 3 post infection. Mice were also treated with 4 μg mIFNβ neutralising antibodies (Leinco Technologies Inc). Control groups were treated with either buffer or an IgG isotype control antibody equivalently.

### Plaque Assay – detection of infectious virus and collection of vaginal wash samples

Vaginal washes were collected by pipetting 30 μL of PBS into the vaginal cavity. Washes were performed twice for each mouse and pooled. Samples were immediately transferred to dry ice and stored at −80°C until ready for use. Then Vero cells in 12-well trays at approximately 90% confluency were infected with 300 μL of serially-diluted virus-containing supernatants for 1 hour at 37 °C. Supernatants were then replaced with a 1mL overlay of complete media containing 1.5% (w/v) carboxymethylcellulose (CMC) (Sigma) and cells returned to culture for 5 days. Cell monolayers were then fixed with 10% formalin and incubation for 1 h. The CMC overlay was then removed, and plaques were visualised via crystal violet stain. Plaques were enumerated, and virus infectivity expressed as plaque-forming units (PFU) per mL.

### Tissue RNA extractions and cDNA preparation and qRT-PCR

500 μL of TRISURE reagent was added to each frozen tissue and these were dissociated in an RNAse free microcentrifuge tube (Eppendorf) using a homogenisation pestle. A further 500μL of TRISURE was added prior to the addition of 200 μL chloroform and separation of the aqueous phase by cold centrifugation. The aqueous layer was reserved and 500 μL 100% ethanol was added. This was then transferred to an RNAeasy column (QIAGEN) for final RNA isolation as per manufacturer’s instructions. cDNA synthesis was performed using MMLV-reverse transcriptase (Promega), reactions contained either 500 ng or 1000 ng RNA (dependent on the lowest RNA yield in a tissue cohort for consistency) and primed with 500ng random hexamer. cDNA were diluted 1 in 4 prior to use. qRT-PCR was performed with Roche FastStart Universal SYBR green (ROX) reagent using the QuantStudio 7 Real-Time PCR System (Applied Biosystem).

### Quantification and analysis of viral RNA (vRNA) genome copies per μg RNA determination of lower LOD for uninfected tissues

A plasmid containing the full-length genome of a Brazilian ZIKV (Paraiba_2015) was used to generate a standard curve for qRT-PCR analysis [58]. First the plasmid was linearised using EcoRV and column purified. The number of genome copies per ng of plasmid was enumerated based on the molecular weight of the plasmid. Serial dilutions on a log2 scale were used to place the curve within a detectable range (between 13 – 36 cycles). A standard curve containing 8 dilutions (including blank) was run with each tissue cohort. Raw CT values were plotted against log2 transformed genome copies and CT values from each tissue cohort were interpolated using the standard curve. Log2 interpolated values were then transformed to a linear scale and normalised to μg RNA used in total cDNA preparation. To achieve normal distributions and equal standard deviations between genotypes, data was log10 transformed and represented as log10 genome copies per μg RNA. Determination of the limit of detection for uninfected tissues was performed by qRT-PCR analysis of treatment, genotype and tissue matched uninfected samples (5 per group) using the same ZIKV PCR primers. CT values from uninfected tissues were compared between genotypes and tissues to ensure consistency in primer background. The earliest detected CT value for all uninfected tissues was determined to be 36.4. This value was then applied to each tissues standard curve to give the minimum value for infected tissue detection. Mice were determined to be uninfected by assessment of melt curves in both technical duplicates and their CT value compared to uninfected controls.

### Viral RNA In-Situ hybridization

RNA ISH was performed on formalin fixed paraffin embedded sections (5 μM) using RNAscope 2.5 HD (Brown) (Advanced Cell Diagnostics) according to the manufacturer’s instructions and as previously described [59].

### RNA-seq sample preparation

Ect1 and VK2 cells seeded to 90% confluency in 6 well plates were treated with hlFNε, hlFNα-2a or hIFNλ-3 (100 ng/mL) diluted in fresh culture media or left untreated (n = 4) for 6 h prior to harvesting RNA in RLT buffer (Qiagen) with added β-ME. Lysates were homogenized by passing through a 20-gauge needle attached to a sterile plastic syringe. Total RNA was extracted from lysates using the Qiagen RNeasy kit, including removal of DNA contamination using the Qiagen RNase free DNase set as per the manufacturer’s instructions.

### RNA-seq library preparation

Libraries were generated using an in-house multiplex RNA-seq method (MHTP, Medical Genomics Facility) and were prepared using 20 ng of total RNA input. An 8 bp sample index and a 10 bp unique molecular identifier (UMI) were added during initial poly(A) priming and pooled samples were amplified using a template-switching oligonucleotide. The Illumina P5 and P7 sequences were added by PCR and Nextera transposase, respectively. The library was designed so that the forward read (R1) utilised a custom primer (5′ GCC TGT CCG CGG AAG CAG TGG TAT CAA CGC AGA GTA C 3′) to sequence directly into the index and then the 10 bp UMI. The reverse read (R2) used the standard R2 primer to sequence the cDNA in the sense direction for transcript identification. Paired-end sequencing (R1 19 bp; R2 72 bp) was performed on the NextSeq 550 (Illumina), using the v2.5 High Output Kit.

### RNA-seq analysis

RNA-seq analysis was performed in R (v3.5.1 and v3.6.1). The scPipe package (v1.2.1) [60] was employed to process and de-multiplex the data. A combined FASTQ file was created from the R1 and R2 FASTQ files, by trimming the sample index and UMI sequences and storing them in the read header, using the sc_trim_barcode function (with bs2 = 0, bl2 = 8, us = 8, ul = 10).

Read alignment was performed using the RSubread package (v1.30.9) [61]. An index was built using the Ensembl *Homo sapiens* GRCh38 primary assembly genome file and alignment was performed with default settings. Aligned reads were mapped to exons using the sc_exon_mapping function with the Ensembl *Homo sapiens* GRCh38 v98 GFF3 genome annotation file.

The resulting BAM file was de-multiplexed and reads mapping to exons were associated with each individual sample using the sc_demultiplex function, taking the UMI into account, and an overall count for each gene for each was sample was generated using the sc_gene_counting function.

Additional gene annotation was obtained using the biomaRt package (v2.36.1) and a DGEList object was created with the counts and gene annotation using the edgeR package (v3.28.1) [62]. The filterByExpr function was used to remove lowly expressed genes and normalisation factors were calculated using the TMM method.

Samples were assigned into groups and counts were transformed using the voom function and a linear model was fit using the lmFit function from the limma package (v3.42.0) [63]. For each cell line, IFN-treated groups were compared to the respective untreated control, using the contrasts.fit function. Moderated *t*-statistics were calculated using the treat function with a 1.2-fold cut-off. Differentially expressed genes were determined using an adjusted *p*-value < 0.05.

Relative log_2_ counts per million (CPM) expression values were used for heat maps. The edgeR cpm function was used to obtain log_2_ CPM values. Relative values were obtained for each cell line by subtracting the average log_2_ CPM value for the respective untreated samples, obtained using the cpmByGroup function. Heat map scales were truncated to ±4. The HALLMARK_INTERFERON_ALPHA_RESPONSE gene set was extracted from the from the Broad Institute Molecular Signature Database (v5.2) Hallmark gene set collection [64] using the EGSEA package (v1.14.0).

The original multiplexed R1 and R2 FASTQ files were de-multiplexed using cutadapt v3.0 (with error rate 0 and action none) and uploaded to GEO, along with UMI counts generated by scPipe, with accession number GSE (to be generated upon submission).

### Focus Forming Assay – detection of infectious virus

Vero or Huh 7.5 cells were seeded in 96 well plates (90% confluence for infection) were inoculated with 40 μL of serially diluted viral supernatants for 3 h at 37 °C with manual agitation every 20 minutes. Inoculum was removed and replaced with 100 μL fresh culture medium for 48 h in a level incubator without shaking. Cells were fixed with acetone/methanol prior to indirect immunofluorescence detection of ZIKV protein using the 4G2-panflavivirus E primary antibody and fluorescently labelled secondary antibody. Foci were enumerated by observation under a fluorescent microscope and were defined as more than 3 infected cells in a distinct cluster. Each sample was calculated based on biological triplicates with technical duplicates then expressed as Focus Forming Units (FFU) per mL.

### Plasmids and transfections

Non-structural proteins were PCR amplified from an infectious clone of a Brazilian isolate of ZIKV (Paraiba_2015) [58] and cloned into the expression plasmid pCDNA6.2 by Gibson Assembly using primers containing 3′ FLAG extension. Empty vector was used as a control in all expression experiments or as a filler for co-transfection in dose dependent assays.

NS1

F ACCGATCCAGCCTCCGGACTCTAGAatggatgtggggtgctc

R TCAGTTAGCCTCCCCCGTTTAAACTTACTTGTCGTCATCGTCTTTGTAGTCtgcagtcaccactg

NS2B/3

F ACCGATCCAGCCTCCGGACTCTAGAatgagctggcccccta

RTCAGTTAGCCTCCCCCGTTTAAACTTACTTGTCGTCATCGTCTTTGTAGTCtcttttcccagcgg

NS4A

F ACCGATCCAGCCTCCGGACTCTAGAatgggagcggcttttgg

R TCAGTTAGCCTCCCCCGTTTAAACTTACTTGTCGTCATCGTCTTTGTAGTCtctttgcttttctggctca

NS5

F GGG ACCG ATCCAGCCTCCGG ACTCTAGAATGgggggtggaacag

R GTTTCAGTTAGCCTCCCCCGTTTAAACTTACTTGTCGTCATCGTCTTTGTAGTCcagcactccaggtg

Plasmids containing constitutively active RIG-I-N or ISRE-Luciferase have been described previously [10, 43]. Transfections were performed using with a 2:1 ratio of lipofectamine 2000 reagent (Invitrogen) to DNA.

### Dual luciferase assay

HeLa cells were co-transfected with the ISRE-luciferase reporter construct, the constitutive renilla luciferase reporter pRL-TK (Promega) as a transfection control and the according pCDNA6.2-NS expression construct or empty vector (total DNA lμg per 12 well). Each transfection condition was carried out in triplicate. 24 h following stimulation with IFN cells were harvested in 1 × passive lysis buffer (Promega) and stored at −20°C. For luciferase assays, the samples were thawed and aliquoted in technical duplicates for detection of firefly and renilla luciferase activity using the Dual-Luciferase Reporter Assay kit using the GloMax luminometer to the manufacturer’s specifications (Promega). Promoter activity was determined by normalising firefly luciferase values to renilla luciferase values and expressed as relative light units (RLU).

### Immunofluorescence microscopy

Briefly, cells were grown in culture plates and fixed with ice-cold acetone:methanol (1:1) for 10 min at 4 °C. After washing twice with PBS, samples were blocked with 5% BSA in PBS for 30 min at room temperature and incubated with primary antibody (for specific usage see Sup.) diluted in PBS/1% BSA for 1 h at room temperature. After washing twice with PBS, cells were incubated with Alexa Fluor-conjugated secondary antibody diluted 1:200 in PBS/1%BSA for 1 h at 4 °C in the dark. Samples were then washed with PBS and incubated with DAPI (Sigma-Aldrich, 1 μg/mL) for 5 min at room temperature. Samples were then washed with PBS. Images were then acquired using a Nikon TiE inverted fluorescent microscope. Contrast was applied using the ‘Autoscale’ function of NIS.

### Western blot for STAT1/STAT2 phosphorylation

HeLa cells were seeded in 6 well trays to be approximately 80 % confluent for transfections. Transfections of NS protein expression constructs or empty vector control were performed as previously described using lipofectamine 2000. 24 h post transfection HeLa cells were stimulated with type-I IFN at the indicated concentrations, 30 minutes later protein was harvested in lysis buffer and left in the well on ice for 10 minutes prior to transfer into a new tube. Lysates were then homogenised by 10 passages through a 25 G needle prior to clearing cellular debris by centrifugation (16,000 RCF for 10 minutes at 4°C). Cleared lysate was diluted with reducing loading dye (3:4) and heated at 95 °C for 5 minutes prior to loading into SDS acrylamide gel and run for 90 minutes. Separate gels were run for each protein of roughly the same molecular weight on the same day to prevent freeze / thaw degradation. Membrane transfer was done for 1 h in cold transfer buffer. Primary antibodies were applied at 1: 1000 (or to the recommended dilution) overnight in 1 % skim milk at 4°C with agitation. The following day secondary HRP conjugated antibodies were applied to the manufacturer’s specification for 1 h at room temperature with agitation. SuperSignal™ West Femto Maximum Sensitivity Substrate (ThermoFisher) was used to detect HRP signal using the BioRad gel dock imager.

### Statistical analyses

For *in vivo* analysis of vaginal washes, the ordinary two-way ANOVA was used on log10 transformed data comparing independent time points. For qRT-PCR analysis of vRNA the ordinary one-way ANOVA was used on log10 transformed data compared to wildtype. All graphing and statistical analyses were performed using GraphPad Prism 8.0. For *in vitro* experiments the analysis method is described in the figure legend.

## Acknowledgements

For conducting ISH experiments we would like to acknowledge Greg Saturday and Rebecca Rosenke (Rocky Mountain Veterinary Branch, RML, NIAID).

## Funding

This research was funded by the National Health and Medical Research Council (NHMRC) of Australia, awarded to M.R.B (APP1145613). Additionally, this work was supported by the Victorian State Government Operational Infrastructure Scheme. M.D.T was supported by an NHMRC Career Development Fellowship (1123319). Funding was also provided in part by the Division of Intramural Research, National Institutes of Health, National Institute of Allergy and Infectious Diseases.

## Author contributions

R.C.C-S, P.J.H and M.R.B conceived the experiments, provided intellectual input and wrote and edited the manuscript. P.J.H and M.R.B secured funding. M.D.T provided intellectual input and performed mice experiments with assistance from R.C.C-S and S.R. L.J.G and J.A.G performed mRNASeq experiments and analysis. R.C.C-S and O.R performed all *in vitro* experinets. S.A.H, B.S, N.S.E and K.H.V provided essential reagents while S.S.L produce in-house recombinant IFNε. S.M.B performed in-situ hybridisation experiments, provided NS5 Ab and intellectual input.

## Notes

### Competing Interest Statement

The authors have declared no competing interest.

