## Supplementary Figures for "Constitutive expression and distinct properties of IFN-epsilon protect the female reproductive tract from Zika virus infection"

### SUPLIMENTARY

### Supplementary 1

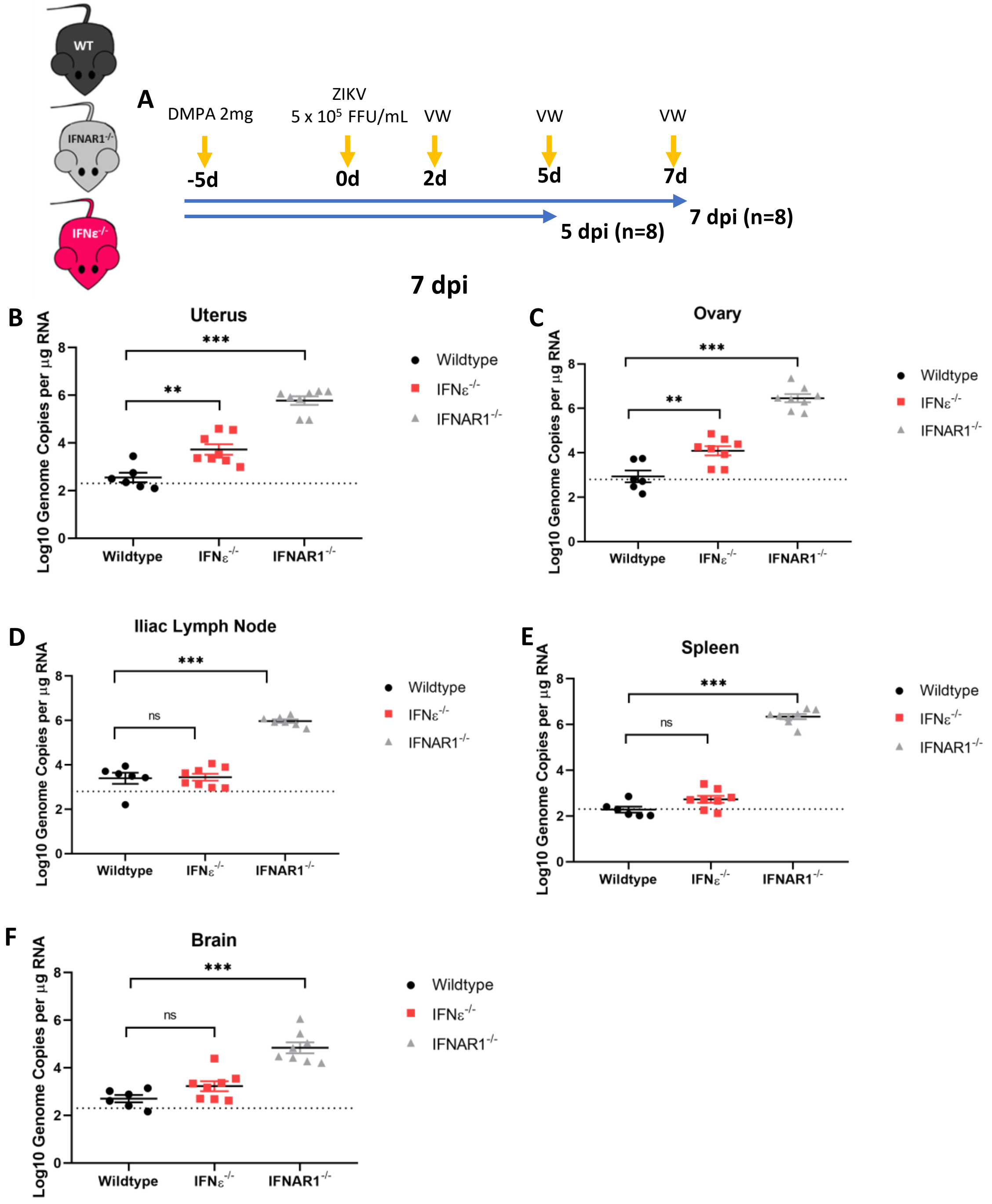


**Sup. Fig 1:** ZIKV replication is inhibited by IFNε in a mouse model of vaginal transmission. A) Experimental time line of WT (black), mice lacking IFNε (IFNε-/-) (red) or mice lacking the type-I IFN receptor (IFNAR1-/-) (grey) were infected with ZIKV at 5 X10^5 FFU 5 days post DMPA treatment and vaginal washes were taken at 2, 5, 7 dpi. 8 mice were culled at 5 dpi and 8 were culled at 7 dpi. B, C, D, E, & F) Tissues taken at 7 dpi were used to harvest RNA for analysis of viral RNA by qRT-PCR in the uterus, ovary, illiac lymph node spleen and brain respectively.

### Supplementary 2

A

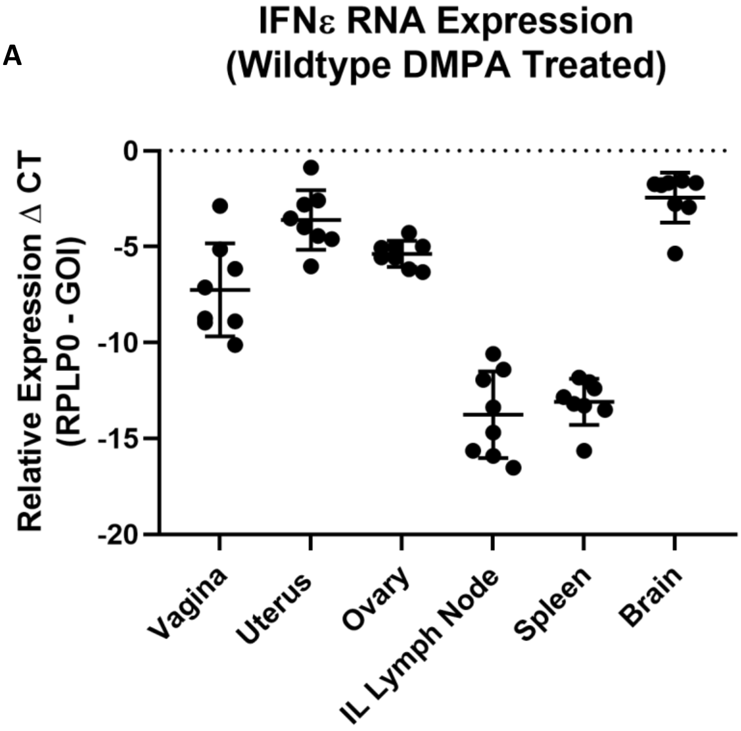


B

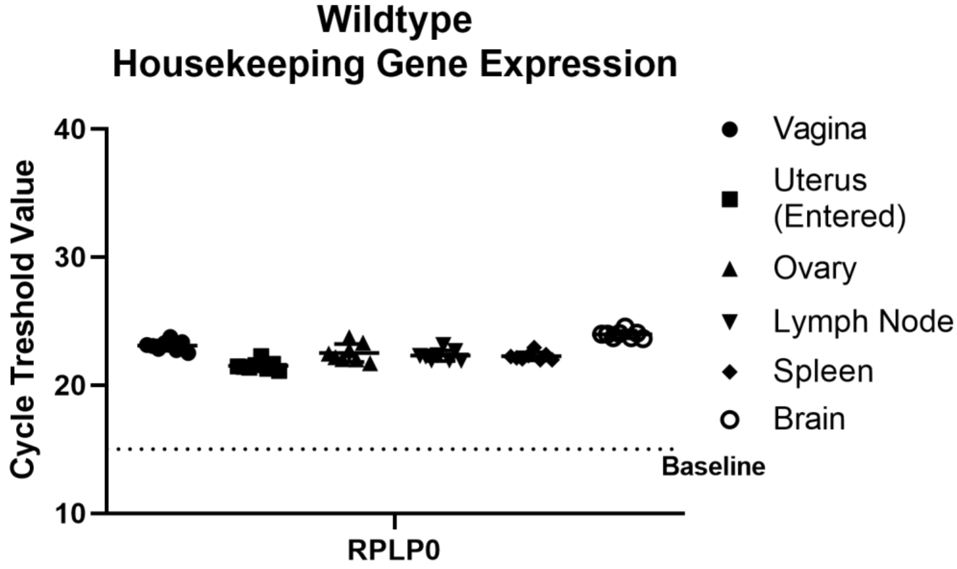


**Sup. Fig 2:** IFNε is highly expressed by tissues of the FRT and in the brain of WT mice. A) Uninfected mice (n=8) were culled 10 days post DMPA treatment to mirror the timeline of the experiment shown in Figure 1 and tissues were collected as previously described. A) the level of IFNε RNA was determined by qRT-PCR and was expressed as ΔCT normalised to expression of the housekeeping gene RPLP0 (36B4). B) Expression of RPLP0 was compared between the tissues by qRT-PCR and expressed as raw cycle threshold value.

### Supplementary 3

**A**

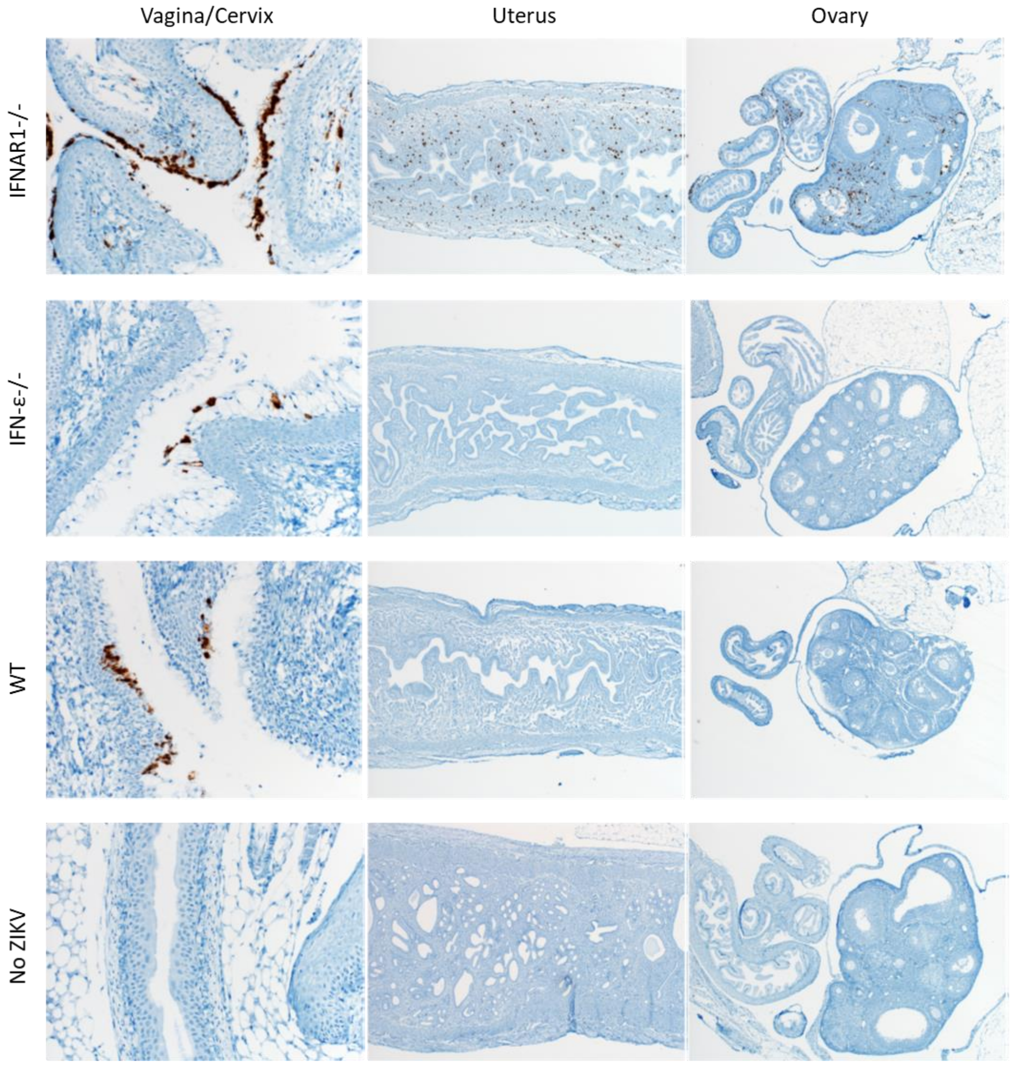


**B**

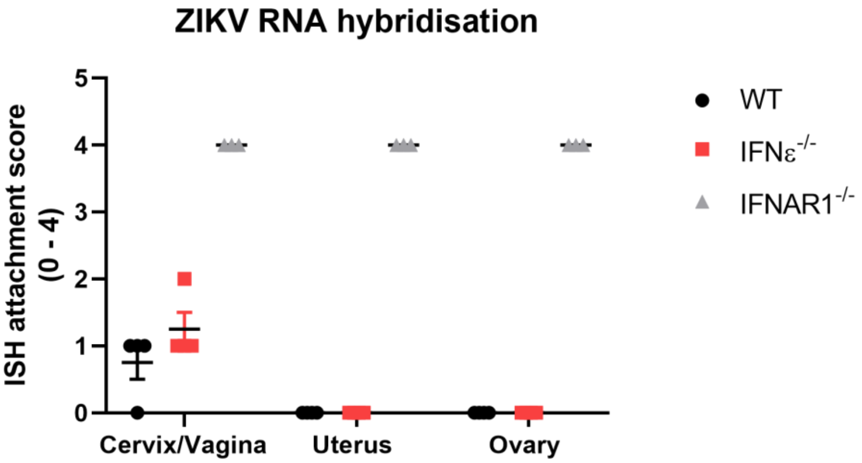


**Sup. Fig 3:** RNAscope in-situ hybridisation (ISH) of ZIKV RNA in FRT tissues at 5 dpi (n = 4)**.** Whole FRT tissues (vagina, cervix, uterus and ovary) were fixed in formalin for 24 h prior to paraffin embedding and sectioning (5 μM), The RNAscrope protocol was used to the manufacturers specifications to detect ZIKV infection using a probe specific to ZIKV –ssRNA. A) Representative ISH images detecting from WT, IFNε-/- and IFNAR-/- mice (ZIKV in brown). B) Pathologist scoring of ISH attachment ( 0 = none, 1 = rare/few, 2 = scattered, 3 = moderate, 4 = numerous).

### Supplementary 4

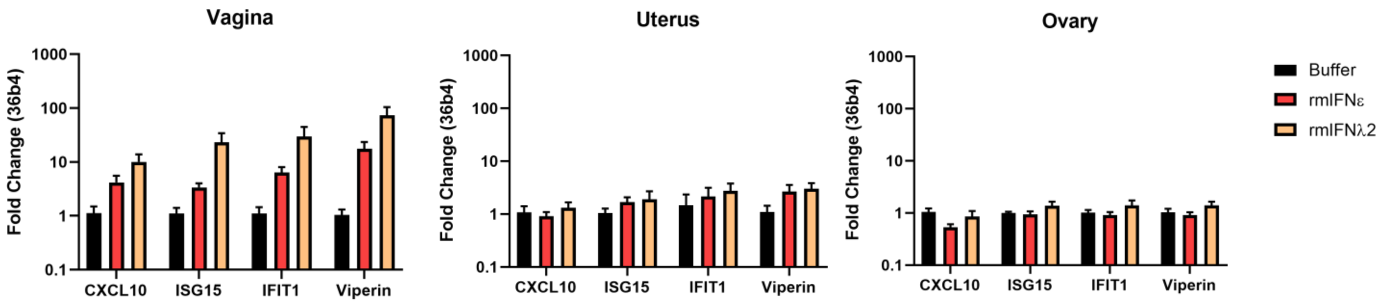


**Sup. Fig. 4: A, B & C)** ISG induction in the vagina, uterus and ovary in DMPA treated, uninfected, IFNε-/- mice following 6 h iVag treatment with the indicated IFN or buffer control.

### Supplementary 5

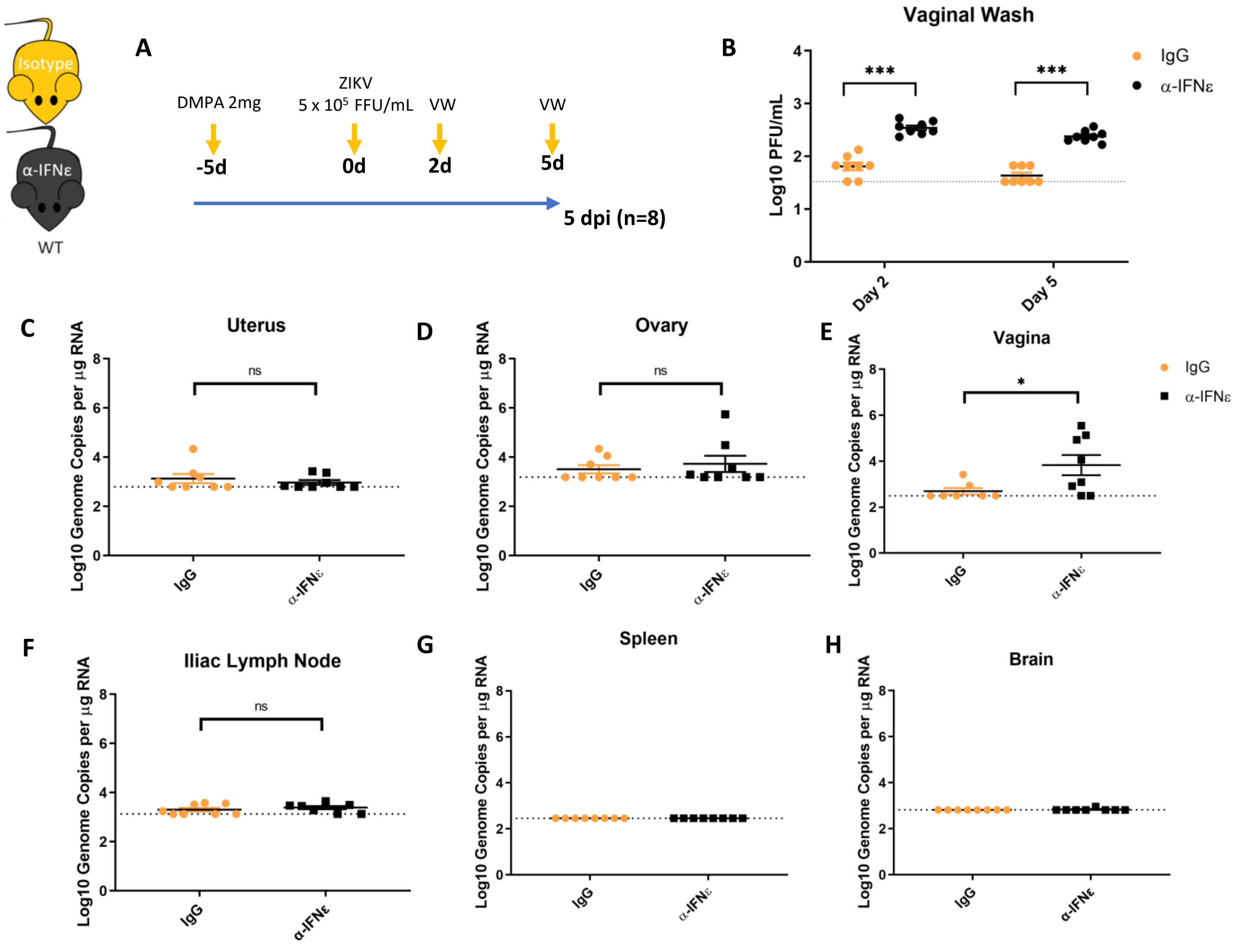


**Sup. Fig. 5:** iVag treatment of wildtype mice with IFNε neutralising antibody enhances ZIKV infection in the vagina (Fig. 1) but not in other tissues. **A)** Experimental time line of (WT mice were treated 6 h prior to infection with 100 ug anti-IFNε or isotype control, then infected with ZIKV at 5 X10^5 FFU 5 days post DMPA treatment and vaginal washes were taken at 2, 5, mice were culled at 5 dpi. **B)** Infectious virus was measured from vaginal washes by plaque assay at 2 and 5 dpi. **C, D, E, F, G)** Tissues taken at 5 dpi were used to harvest RNA for analysis of viral RNA by qRT-PCR in the uterus, ovary, vagina, illiac lymph node spleen and brain respectively.

### Supplementary 6

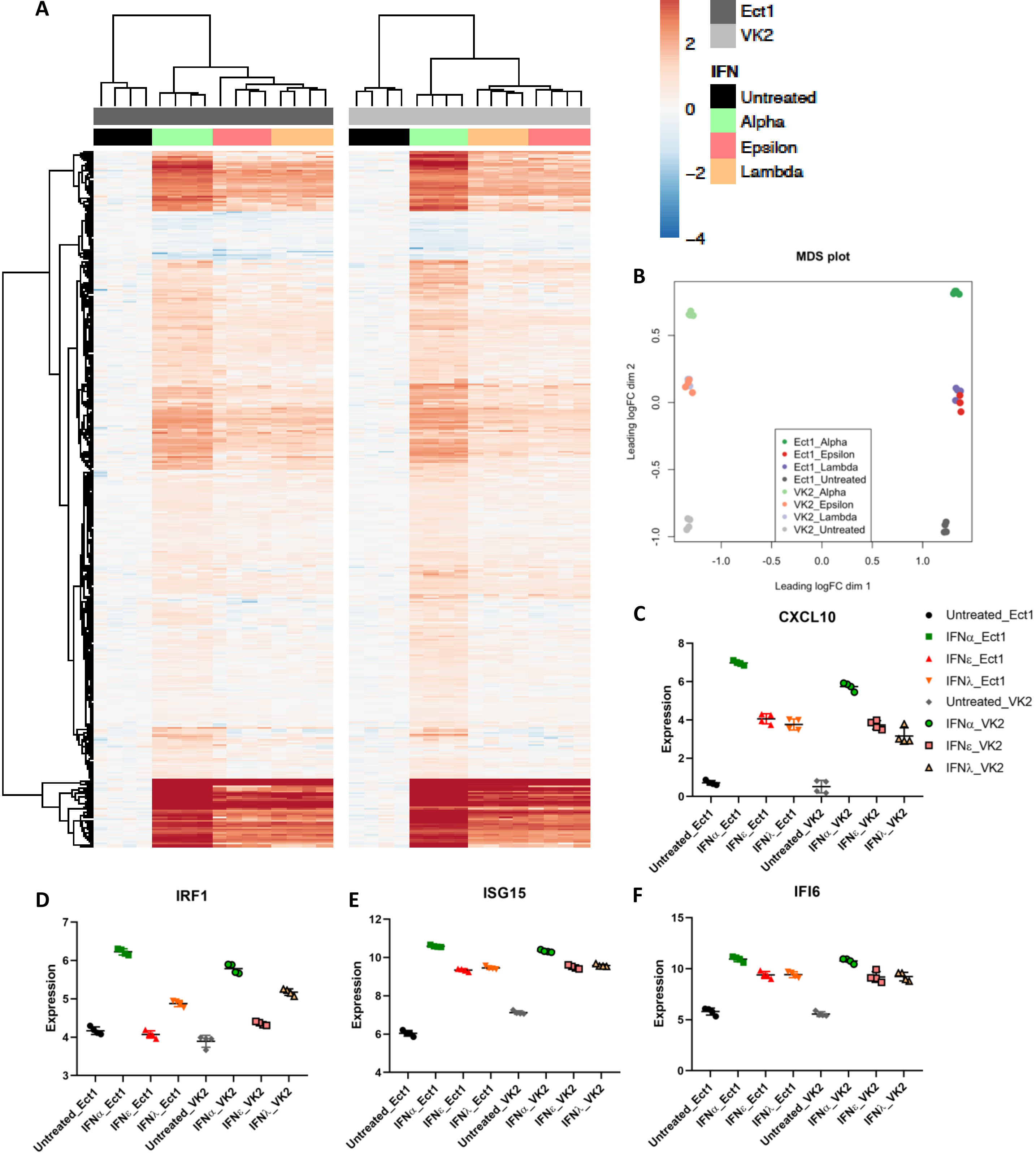


**Sup. Fig. 6:** Ect1 and VK2 cells were treated with IFNε, IFNα-2a or IFNλ-3 (100 ng/mL) or left untreated (n = 4) for 6hr prior to RNASeq analysis (NextSeq550 V2.5). Differentially expressed genes were determined with a 1.2-fold cut-off and adjusted p-value < 0.05. A) Heat map showing expression of all differentially regulated genes. B) MDS plot showing the relationship between samples based on the top 500 most variable genes. C, D, E & F) Expression plots for CXCL10, IRF1, ISG15 and IFI6 respectively in Ect1 or VK2 cells.

### Supplementary 7

**A**

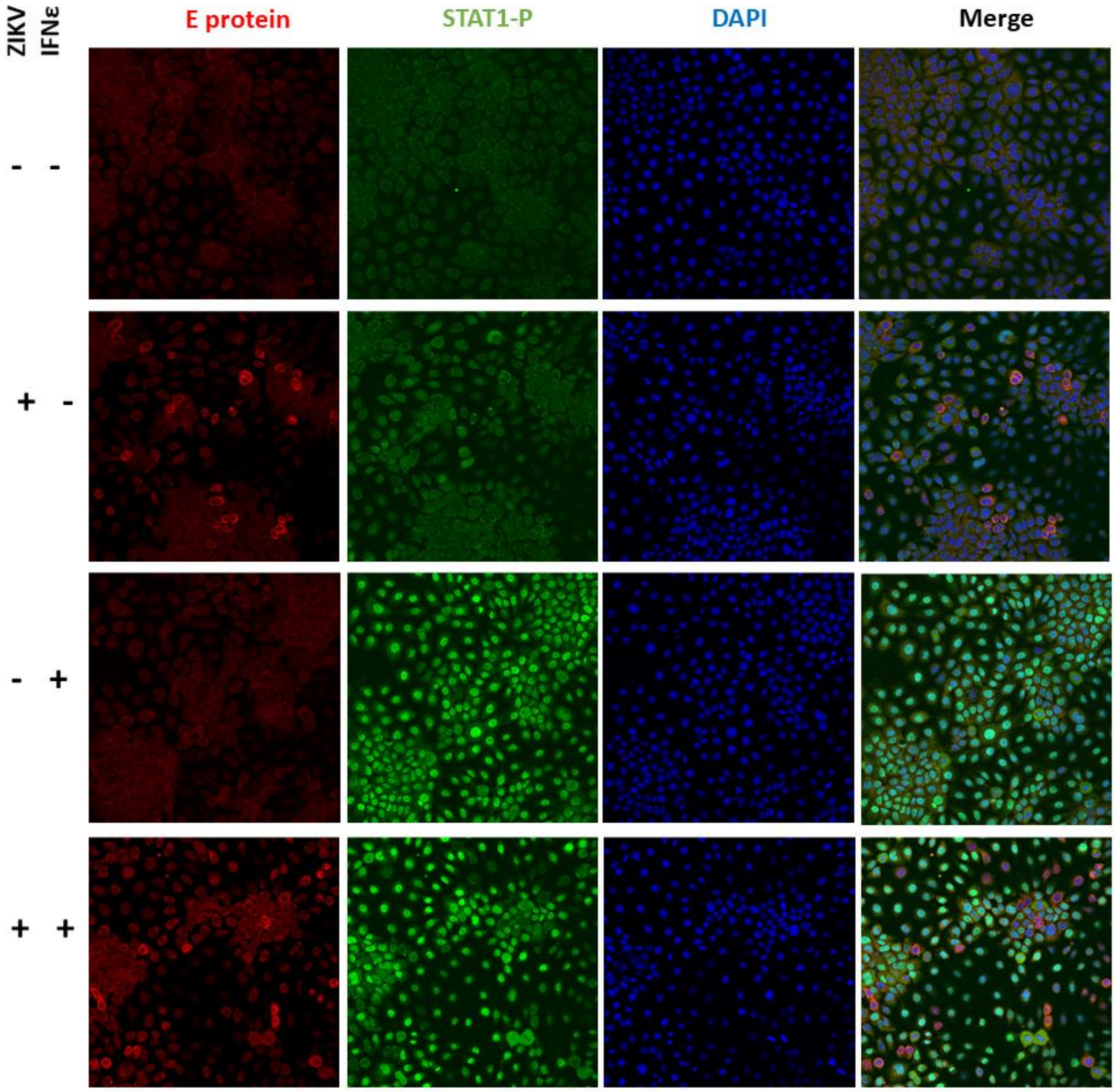

### Supplementary 7

B

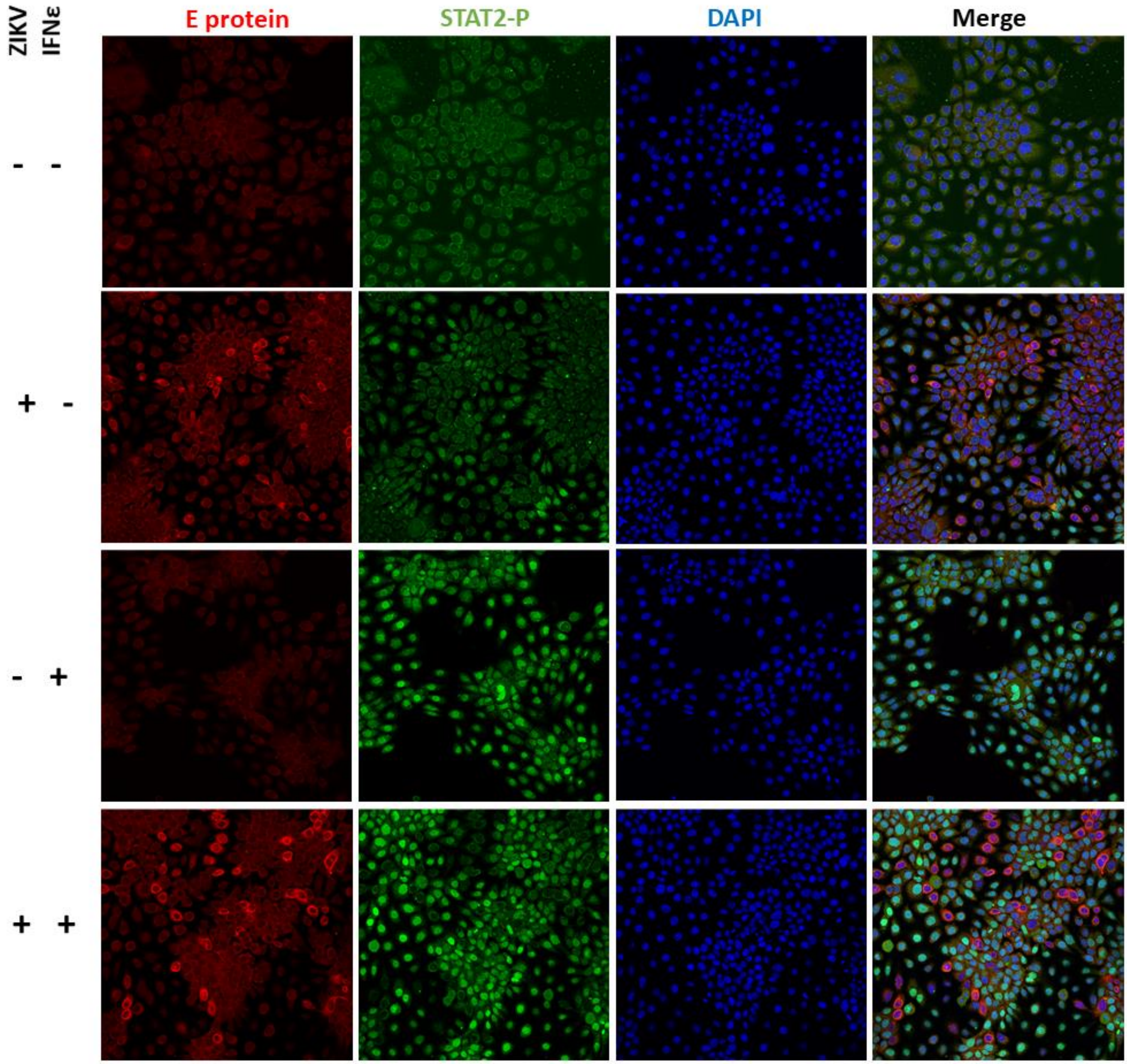

### Supplementary 7

C

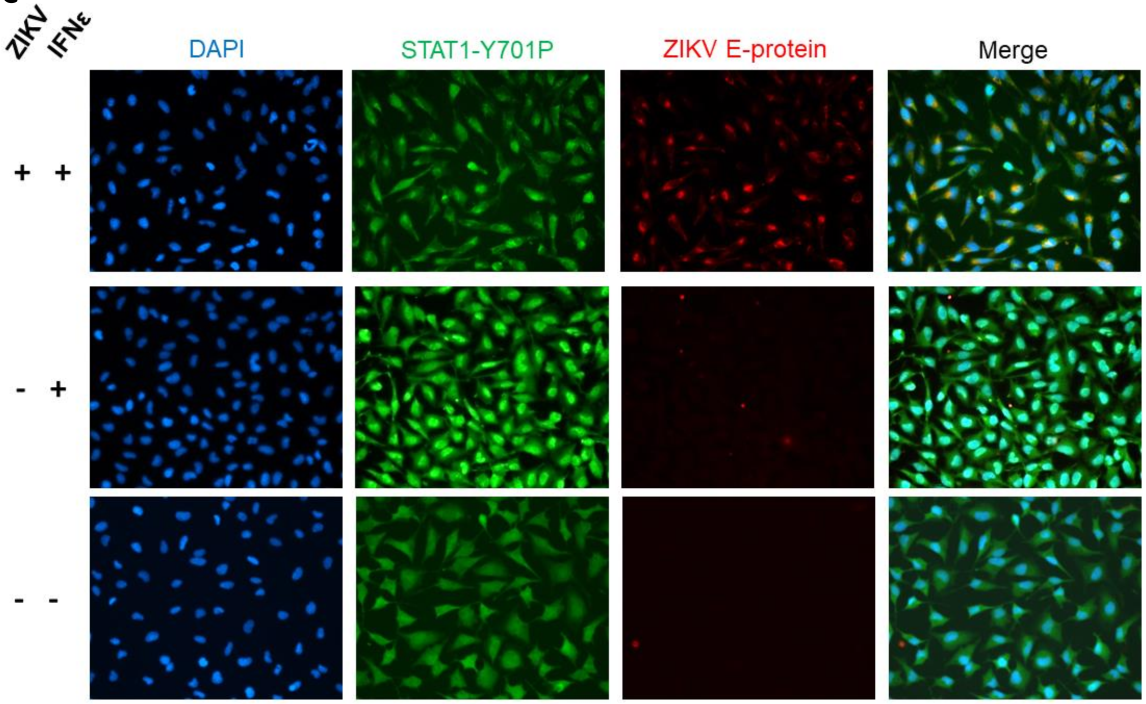


D

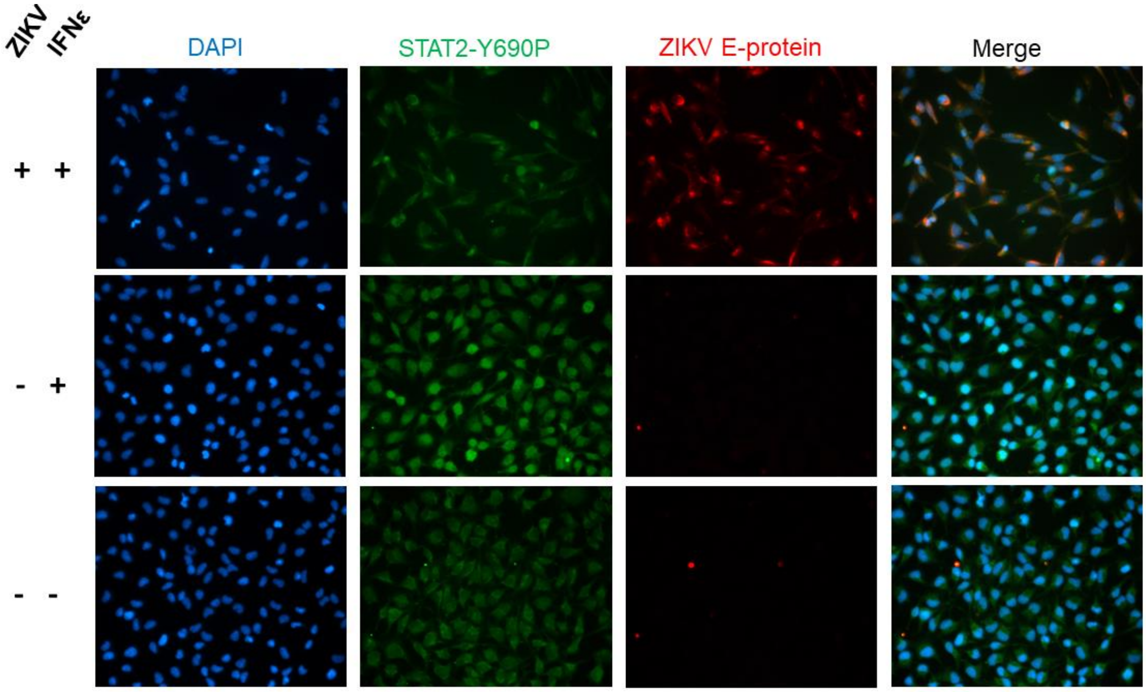


**Sup. Fig. 7: A & B)** Ect1 cells were infected with ZIKV MOI of 10, 24h post infection cells were stimulated with hIFNε (100 U/mL) for 30 min then fixed with acetone/methanol for detection of ZIKV E antigen (Red) and phosphorylated STAT1 or STAT2 proteins (Green) by indirect immunofluorescence, DAPI (Blue). **C & D)** HeLa cells were infected with ZIKV MOI of 10, 24h post infection cells were stimulated with mIFNε (10 U/mL) for 30 min then fixed with acetone/methanol for detection of ZIKV E antigen (Red) and phosphorylated STAT1 or STAT2 proteins (Green) by indirect immunofluorescence, DAPI (Blue).

### Supplementary 8

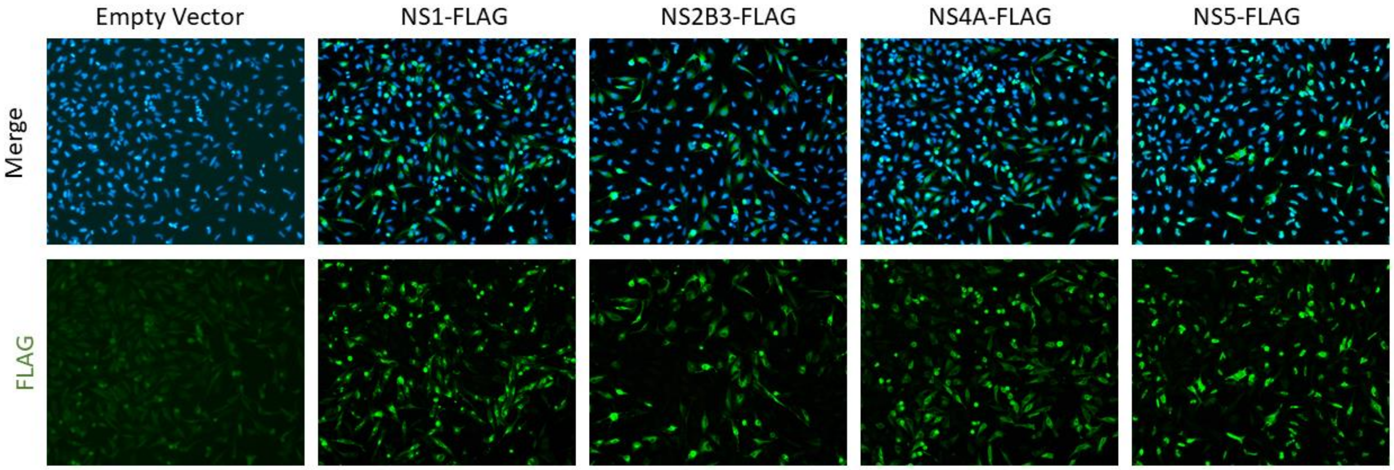


**Sup. Fig. 8:** Transfections of ZIKV NS-FLAG expression constructs were performed in parallel to the experimental data presented in Fig. 7b, indicating the levels of NS protein expression.

### Supplementary 9

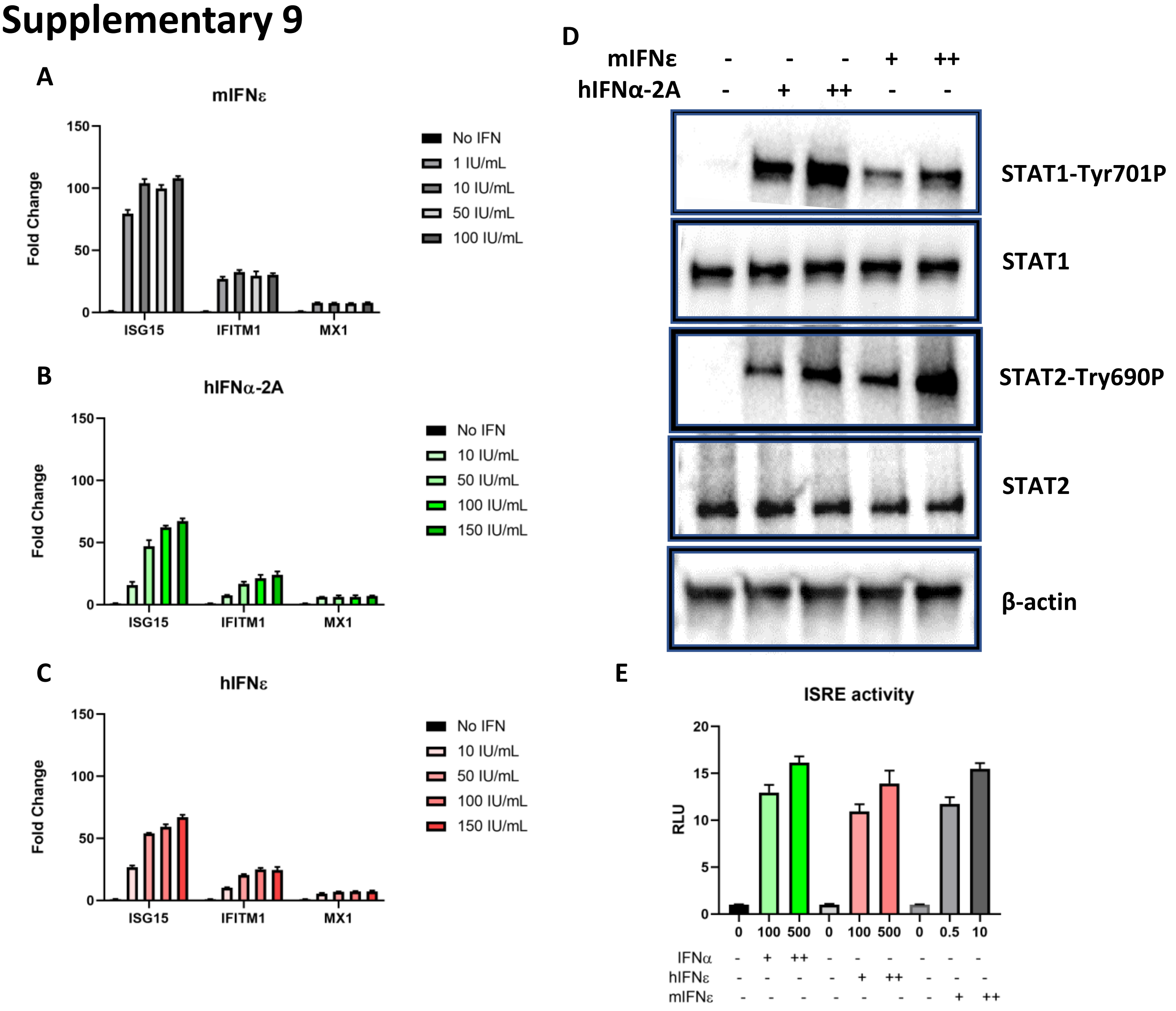


**Supl. Fig. 9:** Murine IFNε stimulate the ISG expression in human cell lines via ISRE driven JAK/STAT pathway activation. **A, B & C)** Dose dependent ISG induction in HTR8 cells following 6h stimulation with indicated amounts of type-I IFNs, data is expressed as fold change relative to untreated cells. **D)** Immunoblot of STAT1/STAT2 phosphorylation in HeLa cells 30 minutes post stimulation with mIFNε (0.5 or 10 U/mL) or hIFNαt-2A(100 or 500 U/mL). **E)** ISRE promoter activity in HeLa cells transfected with an ISRE luciferase reporter was measured by dual luciferase assay in relative light units (RLU) 8 h post stimulation with the indicated amounts of mIFNε.

### Primers

| Gene/Target | Sequence (5’ to 3’) |
|---|---|
| 36B4 (mouse and human) | F – AGA TGC AGC AGA TCC GCA T<br><br>R – GGA TGG CCT TGC GCA |
| Human IFNε | F – TCA GCC TCT TCA GGG CAA ATA<br><br>R – GAG GAA TTT CTC CGT GTG GTT T |
| Human IFN-B (Published) | F – GCA GTC TGC ACC TGA AAA GAT ATT<br><br>R – TGT ACT CCT TGG CCT TGA GGT A |
| Human IFNL1 | F – GGA AGA GTC ACT CAA GCT GAA AAA C<br><br>R – AGA AGC CTC AGG TCC CAA TTC |
| Human IFNL2-3 | F – CAG CTG CAG GTG AGG GA<br><br>R – GCG GTG GCC TCC AGA ACC TT |
| human ISG15 | F – TGG CGG GCA ACG AAT T<br><br>R – GGG TGA TCT GCG CCT TCA |
| human IFIT1 | F – AAC TTA ATG CAG GAA GAA CAT GAC AA<br><br>R – CTG CCA GTC TGC CCA TGT G |
| human Viperin | F – GTG AGC AAT GGA AGC CTG ATC<br><br>R – GCT GTC ACA GGA GAT AGC GA |
| human CXCL10 (IP-10) | F – TCC ACG TGT TGA GAT CAT TGC<br><br>R – TCT TGA TGG CCT TCG ATT CTG |
| human CXCL11 | F – CCT TGG CTG TGA TAT TGT GTG C<br><br>R – CCA CTT TCA CTG CTT TTA CCC C |
| human IRF1 | F – CCA GCC CTG ATA CCT TCT CTG A<br><br>R – AAG TCC TGC ATG TAG CCT GGA A |
| human IFI6 | F – CTG AAG ATT GCT TCT CTT CTC<br><br>R – CAC TTT TTC TTA CCT GCC TC |
| Murine IFNE | F – GAA ACG GAT TCC CTT CCA AT<br><br>R – ACT GCT GGA CTG ACG AGC TT |
| Murine IFNA | F – CTG CCT GAA GGA CAG GAA GG<br><br>R – GTC ATT GAG CTG CTG GTG GA |
| Murine IFNB | F – AGA AAG GAC GAA CAT TCG GAA A<br><br>R – CCG TCA TCT CCA TAG GGA TCT T |
| Murine IFNL2 | F – CCA CAT TGC TCA GTT CAA GTC TCT<br><br>R – TCC TTC TCA AGC AGC CTC TTC T |
| murine ISG15 | F – GGG GCC ACA GCA ACA TCT AT<br><br>R – AGC CAG AAC TGG TCT TCG TG |
| murine IFIT1 | F – TGG CGT AGA CAA AGC TCT TCA TC<br><br>R – TAG CAG AGC CCT TTT TGA TAA TGT AA |
| murine Viperin | F – TTG GGC AAG CTT GTG AGA TTC<br><br>R – TGA ACC ATC TCT CCT GGA TAA GG |
| murine HPRT | F – AAG CTT GCT GGT GAA AAG GA<br><br>R – TTG CGC TCA TCT TAG GCT TT |
| murine CXCL10 | F – ATG ACG GGC CAG TGA GAA TG<br><br>R – ATG ATC TCA ACA CGT GGG CA |
| ZIKV PRVABC59 – prM specific | F – GTG TGA TGC CAC CAT GAG CTA<br><br>R – TGG CAG GTT CCG TAC ACA AAC |

### Antibodies

| Antibody | Usage | Primary (1º) or Secondary (2º) | Dilution | Incubation | Supplier |
|---|---|---|---|---|---|
| Mouse anti - flavivirus E (4G2) hybridoma supernatant | Immunofluorescence | 1º | 1/5 | RT 1h | Made in house from hybridoma HB-112 |
| Mouse anti – FLAG | Immunofluorescence | 1º | 1:200 | RT 1h | Sigma Aldrich |
| Rabbit anti – STAT2-Y690P (#D3P2P) | Immunofluorescence | 1º | 1:100 | 4 ºC overnight | Cell Signaling |
| Rabbit anti – STAT1-Y701P (#58D6) | Immunofluorescence | 1º | 1:100 | 4 ºC overnight | Cell Signaling |
| Goat anti-Mouse IgG, Alexa Fluor 555 linked | Immunofluorescence | 2º | 1:200 | On ice 1h | Invitrogen |
| Goat anti-Mouse IgG, Alexa Fluor 488 linked | Immunofluorescence | 2º | 1:200 | On ice 1h | Invitrogen |
| Rabbit anti – STAT2-Y690P (#D3P2P) | Western Blot | 1º | 1:1000 | 4 ºC overnight | Cell Signaling |
| Chicken anti – NS5 | Western Blot | 1º | 1:1000 | 4 ºC overnight | Sonja Best, Rocky Mountains, NIH |
| Mouse anti – FLAG | Western Blot | 1º | 1:1000 | 4 ºC overnight | Sigma Aldrich |
| Mouse anti – βactin | Western Blot | 1º | 1:10000 | 4 ºC overnight | Sigma Aldrich |
| Rabbit anti – STAT1-Y701P (#58D6) | Western Blot | 1º | 1:1000 | 4 ºC overnight | Cell Signaling |
| Rabbit anti – STAT2 (#D9JL) | Western Blot | 1º | 1:1000 | 4 ºC overnight | Cell Signaling |
| Rabbit anti – STAT1 (#D1K9Y) | Western Blot | 1º | 1:1000 | 4 ºC overnight | Cell Signaling |
| Goat anti-mouse IgG (H+L), HRP linked | Western Blot | 2º | 1:10000 | RT 1h | Invitrogen |
| Goat anti-rabbit IgG, HRP linked (#7074) | Western Blot | 2º | 1:1000 | RT 1h | Cell Signaling |
